# Discovering Biomarker Proteins and Peptides for Parkinson’s Disease Prognosis Prediction with Machine Learning and Interpretability Methods

**DOI:** 10.1101/2023.05.18.541380

**Authors:** Ho-min Park, Espoir Kabanga, Dongin Moon, Minjae Chung, Jiwon Im, Yujin Kim, Arnout Van Messem, Wesley De Neve

**Author notes:** these authors contributed equally to this work (Dongin Moon, Minjae Chung, Jiwon Im, Yujin Kim).

## Abstract

Parkinson’s disease is a neurodegenerative disorder that affects millions of people worldwide, posing significant challenges for diagnosis and treatment. This study presents a machine learning pipeline for identifying candidate biomarker proteins and peptides from cerebrospinal fluid mass spectrometry (CSF-MS) tests in Parkinson’s disease patients. Our pipeline comprises two main stages: (1) model training using mutual information-based feature selection and five different machine learning regressors and (2) identification of candidate biomarkers by combining three types of interpretability methods. Our regression models demonstrated promising effectiveness in predicting the Movement Disorder Society-Unified Parkinson’s Disease Rating Scale (MDS-UPDRS) scores, with UPDRS-1 receiving the best predictions, followed by UPDRS-3 and UPDRS-2. Furthermore, our pipeline identified 11 proteins and peptides as potential biomarkers for Parkinson’s disease, excluding Levodopa usage which trivially has the most significant impact on the prognosis prediction. Comparisons with four additional pipelines confirmed the effectiveness of our approach in terms of both model performance and biomarker identification. In conclusion, our study presents a comprehensive machine learning pipeline that demonstrates effectiveness in predicting the severity of Parkinson’s disease using CSF-MS tests. Our approach also identifies potential biomarkers, which could aid in the development of new diagnostic tools and treatments for patients with Parkinson’s disease.

## Introduction

According to the World Health Organization (WHO), Parkinson’s disease is a degenerative condition of the brain, leading to cognitive impairment, mental health disorders, sleep disorders, pain, and sensory disturbances [1]. As pointed out in a technical brief issued by the WHO in 2022, the prevalence of Parkinson’s disease reached an estimated 8.5 million individuals in 2019. In the United States, a recent study supported by the Parkinson’s disease Foundation found that approximately 90,000 individuals are diagnosed with Parkinson’s disease annually, indicating a substantial 50% rise from the previously projected rate of 60,000 diagnoses per year. It should be clear that the incidence and prevalence of this disease have rapidly increased in the past two decades [2]. Clinically, Parkinson’s disease presents with motor symptoms including tremor (a neurological condition characterized by rhythmic oscillatory movements in one or multiple body regions, typically affecting the hands), rigidity, bradykinesia, postural instability, and gait alterations [3]. These symptoms are quantified by converting them into the four parts of the Movement Disorder Society-Unified Parkinson’s Disease Rating Scale (MDS-UPDRS) scoring system [4] during hospital visits, and treatment methods are determined based on the resulting scores.

Various statistical and machine learning approaches have been introduced to identify the causes and symptoms of Parkinson’s disease, leveraging different types of information, including cerebrospinal fluid mass spectrometry (CSF-MS) data and mobility observations. CSF-MS is a diagnostic test used to analyze the proteins in the cerebrospinal fluid to detect neurological disorders. This method has made a significant contribution to the discovery of *α*-synuclein, a major biomarker of Parkinson’s disease, and is still widely used as a tool for diagnosing and tracking Parkinson’s disease [5, 6]. Given that CSF-MS data contains information about hundreds to thousands of proteins and peptides, comprehensive analysis by humans presents a challenge. Therefore, many studies have proposed the use of machine learning methods to analyze CSF-MS data. [7, 8, 9], for example, used machine learning methods to classify individuals with Parkinson’s disease and healthy individuals using CSF-MS data. Based on the fact that Parkinson’s disease causes problems with muscle use, mobility observations such as voice, gait, and finger writing were used to train machine learning models to learn the difference between patients and healthy people [10, 11, 12, 13].

Additionally, clinical datasets such as body mass and cholesterol [14], single photon emission computed tomography (SPECT) images [15], and genetic information datasets consisting of RNA sequences [14, 16] were used to train machine learning models. Compared to mobility and other categories, CSF-MS targets internal proteins in the body, making it the closest approach to identifying the cause, rather than just the symptoms. This method also has the potential to be used to identify signs of this incurable disease in advance or to develop new drugs.

In recent years, the need for interpretable machine learning models has grown significantly, as they facilitate understanding and trust in model predictions, particularly in high-stakes applications such as healthcare, finance, and criminal justice [17]. Furthermore, the General Data Protection Regulation (GDPR) issued by the EU emphasizes the “right to explanation,” which grants individuals the right to comprehend the logic behind automated decisions made by algorithms, especially in instances where these decisions significantly impact their lives [18]. In contemporary research, specifically on methods combining machine learning and CSF-MS, an increasing trend towards adopting and integrating these interpretable methods is evident, aiming to promote transparency and foster a better understanding of model behavior [14, 19].

In this study, we propose a machine learning pipeline for identifying candidate biomarker proteins and peptides from CSF-MS tests in Parkinson patients. The pipeline is divided into two main stages: first, model training using mutual information, a feature selection technique, and five different machine learning regressors; secondly, finding candidate biomarkers by combining three types of interpretability methods. Our study distinguishes itself from prior research in the following ways:

1. **Regression problem-solving**: Most existing studies perform classification [5, 9, 19, 20] by comparing data from patients with Parkinson’s disease with data from healthy individuals. In contrast, we aim to solve a regression problem that predicts MDS-UPDRS scores using only the CSF-MS information of patients having Parkinson’s disease. Although interpreting the results as intuitively as classification accuracy is challenging, regression offers the advantage of serving as a prognostic predictor if a time difference exists between input and output values.
2. **Mutual information-based feature selection**: For problems with hundreds to thousands of input dimensions, such as CSF-MS, it is necessary to reduce the number of input dimensions through methods for feature selection or extraction. In this study, we perform feature selection using mutual information (MI). MI is a statistical concept measuring the degree of dependence between two random variables. Various feature selection methods using this concept have been introduced [21, 22, 23].
3. **Combination of three types of interpretability methods**: Related research that leverages interpretability methods [14, 19] investigates a trained model using only one method. However, the use of one interpretability method may be biased, given hypothetical assumptions and the particular design of the method. To overcome this kind of limitations, we obtain the importance ranks of three interpretability methods, using the average rank to identify candidate key proteins and peptides.

The remainder of this paper is organized as follows. In Section Results, we detail the outcomes obtained by the proposed machine learning pipeline, including the identification of candidate biomarker proteins and peptides, as well as the effectiveness of our regression models. In Section Discussion, we look into the interpretation of our results, paying attention to the implications of our findings. Additionally, we also discuss the advantages and limitations of our approach, highlighting potential areas for future research. Finally, in Section Methods, we describe the methodologies employed in this study, covering the MI-based feature selection process, the various machine learning regressors, and the combination of three types of interpretability methods.

## Methods

This section describes all pipelines including our own pipeline which is used for building regressors for obtaining candidate biomarkers (features) from input CSV datasets. To that end, we mainly utilized Python, scikit-learn [24], and Pandas [25]. To guarantee a high level of usability and reproducibility, we also provide our pipeline in the form of a Jupyter notebook (Google Colab), as available at the [Supplementary material (4)](https://colab.research.google.com/drive/1zH3EqbLTioKL4G_TGM0X0U4tBKY1i3e5?usp=sharing). Readers can thus run our code and modify it for their own purposes.

### Problem definition and dataset

In this problem, we are concerned with patients diagnosed with Parkinson’s disease where these patients receive either semi-annual or annual CSF-MS together with MDS-UPDRS. CSF-MS measures the abundance of protein and peptides in the patient – in this case, 227 protein and 971 peptide candidates – whereas MDS-UPDRS [4] is a prognosis score consisting of four values. The dataset utilized in this study is obtained from a Kaggle competition [26] on progression prediction for Parkinson’s disease. This competition involved 248 patients who visited a hospital approximately every six months, leading to the assignment of MDS-UPDRS scores and the semi-annual or annual creation of CSF-MS test results. The objective of the competition was to predict the MDS-UPDRS score after 0, 6, 12, and 24 months using machine learning based on the provided data. Since MDS-UPDRS comprises four scores, the Kaggle competition deals with time series’ prediction and multivariate regression. Symmetric Mean Absolute Percentage Error (SMAPE) is used as an evaluation metric. As this is a regression problem, all 248 patients are considered to have Parkinson’s disease, albeit with varying degrees of progression.

We formulate this problem using four datasets 𝔻^(t)^ with *t* ∈ {0, 6, 12, 24} denoting the timestamp of the aforementioned tests measured in months. Note that the size of each dataset varies with *t* = 6 being considerably smaller than the other ones. A summary of each subset can be found in Table 3a.

Each dataset consists of input-output pairs **x, y ∼** 𝔻^(*t*)^ with **x ∈** ℝ^1195^ and **y ∈** ℝ^4^. In this scenario, *y*_*i*∈{1,2,3,4}_ denotes four different parts of MDS-UPDRS scores. Note that the measurements are taken at four different time stamps (*t* ∈ {0, 6, 12, 24}) and four MDS-UPDRS scores (*i* ∈ {1, 2, 3, 4}). Therefore, we build 16 different machine learning models corresponding to each timestamp-score combination. In this context, we define a machine learning model as *f* (*θ*, **x**) where *θ* denotes the parameters of the model where this model minimizes SMAPE between the predicted values (by the model) and the true targets (*y*_*i*_).

### Pre-processing

During the pre-processing stage, we performed three main procedures on the features: skewness removal, standardization, and missing value imputation. First, we removed the skewness using logarithmic, Box-Cox, and square root transformations. Second, we standardized the features by setting the mean to zero and the standard deviation to one.

Lastly, we classified missing values into Missing Completely at Random (MCAR), Missing at Random (MAR), and Missing not at Random (MNAR). To determine the missingness types, we employed the Student’s t-test to examine the relationship between missingness and other feature values. Based on the results, we filled in the missing values accordingly.

If the number of missing values in a feature is less than 30, we assumed the type of missing value as MCAR since it did not reach the minimum quantity required for the t-test. For the MCAR cases, we imputed the average value of each feature. In contrast, for the MAR and MNAR cases, in which the missing values are related to other feature variables, we utilized the Singular Value Decomposition (SVD) imputation method [27]. SVD imputation fills in missing values by estimating them based on the underlying patterns and relationships in the data, using a matrix factorization technique. By discerning different types of missing values and employing appropriate imputation strategies, our aim was to enhance the accuracy and reliability of subsequent analysis and modeling [28].

### Feature selection

Although CSF-MS should be used as input for the model, the number of feature dimensions (1,195) exceeded the number of data points (1,068), potentially triggering the curse of dimensionality. To address this issue, the MI between CSF-MS and MDS-UPDRS was calculated, and 129 out of 1,195 features (36 proteins and 93 peptides) were selected based on their MI ranking [29]. Additionally, we divided the UPDRS 23b clinical state which describes Levodopa treatments column into “On” (ingested and good prognosis), “Off” (ingested but poor prognosis), and “No” (not ingested or no information) features based on the medication status of a patient, and added them to the list of features using one-hot encoding. In this context, Levodopa is a medicine that greatly affects the prognosis of Parkinson’s disease.

As a result of the feature selection effort described above, the input to machine learning models end up having a feature dimension of 132 (129 CSF-MS features and 3 Levodopa-related features).

### Training and evaluation

For competition, we are required to submit 16 results, corresponding to each timestamp-score combination. For each of those, we train five regression-based machine learning models: linear regression [30], decision tree regression [31], ElasticNet [32], bayesian ridge regression [30], and gradient boosting regression [33]. Each model is trained using five-fold cross-validation and grid search hyperparameter selection [34]. We evaluated each model’s effectiveness using SMAPE, the standard metric used by the underlying Kaggle competition. For each combination, we selected the model with the lowest SMAPE as the final model. Note that the validation or the test datasets are not publicly available, hence the necessity for cross-validation in order to determine the hyperparameter setup and model (one of the five) that leads to the lowest SMAPE score. After determining the best experimental setup, we (re)train the best-performing model with the entire dataset.

### Finding candidate biomarkers

In order to identify the essential features used by the best-performing model, we employ three interpretability methods: built-in feature importance, Permutation Feature Importance (PFI) [35], and Shapley Additive Explanations (SHAP) [36]. These methods provide numerical values. These values represent how significantly the input features contributed to the predictions made by the model that has completed training (feature importance).

The built-in method is an approach to interpretation that relies on the design of the model, and it computes the significance of features through one of the various metrics. For instance, the mean decrease in Gini impurity is employed in a decision tree, while gradient boosting utilizes a reduction in the loss. Additionally, regression families (linear, ridge, ElasticNet) employ coefficients to assess feature importance.

Alas, each method has its limitations. The built-in method depends on the design of the model and may not be universally applicable. PFI estimates can be noisy and sensitive to the specific data used for evaluation, which may lead to unstable feature rankings. SHAP relies on major assumptions, such as feature independence and additivity, which may not hold true for the dataset we use in this study. By using three distinct methods, we aimed to offset the limitations of each individual method for interpretability.

Each interpretability method is individually computed for every time stamp-score combination, and subsequently aggregated. This approach enables us to ascertain a comprehensive importance measure across all time stamp-score combinations. Nevertheless, determining a unified importance measure from the three interpretability methods proved challenging, as each method not only has its unique calculation process but also produces importance values on different scales. To overcome this issue, we assigned rankings to the importance values derived from each of the three methods and then utilized the average ranking to establish the final importance order.

### Checking pipeline performance

To evaluate the performance of our proposed pipeline, we constructed five alternative pipelines and compared their performances to that of our main (proposed) pipeline. Due to the difficulty in comparing the prediction performance of all 16 scores, we focused on UPDRS-1 (*y*_1_) to UPDRS-3 (*y*_3_), excluding UPDRS-4 (*y*_4_) which has poor prediction performance. Then, we averaged the results for UPDRS-1 to 3 in order to generate an average SMAPE score for comparison.

Including the main pipeline, we conduct experiments using six pipelines listed below.

- **Main pipeline**: this is the proposed pipeline which is mainly described in Section Methods.
- **AP-Multivariate**: we used a multivariate prediction model to make four-dimensional predictions simultaneously, rather than predicting each of the four scores individually. This method was expected to improve the learning of the correlation between the input and the entire score. This pipeline followed the same process as our pipeline, but the ML model was replaced with a 3 multivariate prediction model: linear regression, ElasticNet, and decision tree regression.
- **AP-PCA**: we tested the effect of feature extraction using principal component analysis (PCA). The input was reduced from 1,195 dimensions to 10 dimensions using PCA, and the performance was measured using the main pipeline but with only these features.
- **AP-W/O Levodopa**: we trained the model without Levodopa-related features, which are known the greatest impact on helping Parkinson’s disease symptoms. After excluding the three one-hot encoded features from the input, the remaining features were trained using the same process as our pipeline.
- **AP-DNN**: We trained deep neural networks instead of the existing machine learning model. DNNs with 7 different structures in terms of layer depth and width (number of neurons) were created, using the Adam optimizer with a learning rate of 0.001 and a dropout rate of 0.5. The optimal performance of the DNN model was calculated using grid search and 5-fold cross-validation, following the same approach as our pipeline.
- **AP-Selection**: we trained the model without using MI-based feature selection. We tested three cases: (1) when all 227 proteins were used as input, (2) when 965 peptides were used as input, and (3) when all features were used for learning without discarding any of them. The rest of the process was set the same as our main pipeline.

By comparing the Averaged SMAPE values of our main pipeline and the five alternative pipelines, we were able to assess the effectiveness of our feature selection, feature engineering, and machine learning model choices. This comparison provides insight into the importance of each step in our pipeline and guides future improvements to enhance the prediction of Parkinson’s disease progression using CSF-MS data.

## Results

### Training results

Table 3a outlines four distinct datasets, each representing different time stamps, namely 𝔻^(0)^, 𝔻^(6)^, 𝔻^(12)^, and 𝔻^(24)^, with 1,068, 137, 543, and 435 data points respectively. All data points across datasets have 132 features, including the categorical feature of Levodopa treatment status (“No”, “On”, and “Off”). The distribution of Levodopa treatment varies within each dataset, with the highest number of patients not on Levodopa across all datasets.

Table 3b presents the outcomes of our experiment, which are quantified by the SMAPE scores for each pairing of four time stamps and MDS-UPDRS, labelled as *y*_1_, *y*_2_, *y*_3_, and *y*_4_. The datasets denoted as 𝔻^(0)^, 𝔻^(6)^, 𝔻^(12)^, and 𝔻^(24)^ represent distinct time stamps.

The SMAPE scores range from 53.08 to 185.07, indicating the relative accuracy of the model for each timestamp and MDS-UPDRS score combination. SMAPE tends to give a higher penalty for over-predictions, and predicting the fourth UPDRS score, *y*_4_, proved to be very challenging. This can be attributed to the fact that approximately half of the UPDRS-4 scores had missing values, which were filled with a minimum score of 0, thereby leading to a high penalty for over-predicting.

Conversely, the prediction of UPDRS-1 (*y*_1_) was the best, followed by UPDRS-3 (*y*_3_) and UPDRS-2 (*y*_2_), in that order. Notably, the dataset 𝔻^(6)^ showed the lowest SMAPE scores, suggesting the best model performance for this time interval. However, this dataset only contained 137 data points, considerably smaller compared to the other intervals. Therefore, while the model performance on this dataset appears promising, the generalizability of this result is doubtful due to the limited number of data points.

Figure 3c illustrates the quantity and varieties of the models ultimately selected. Of the 16 timestamp-score combinations, eight gradient boosting regression, five decision tree regression, and three ElasticNet were selected as final model. An interesting observation, as can be seen in the [Supplementary material (1)](https://docs.google.com/spreadsheets/d/1lu58P36kbP6XobipBoVMIDs0m656EJ0XLQcO2ZtNIi0/edit?usp=sharing), is that the decision tree was generally the preferred approach for predicting UPDRS-4 (*y*_4_).

### Key proteins and peptides

Table 1 displays the highest-ranked features captured through our pipeline and their associated UniProt ID, description, and average ranking.

**Table 1.**
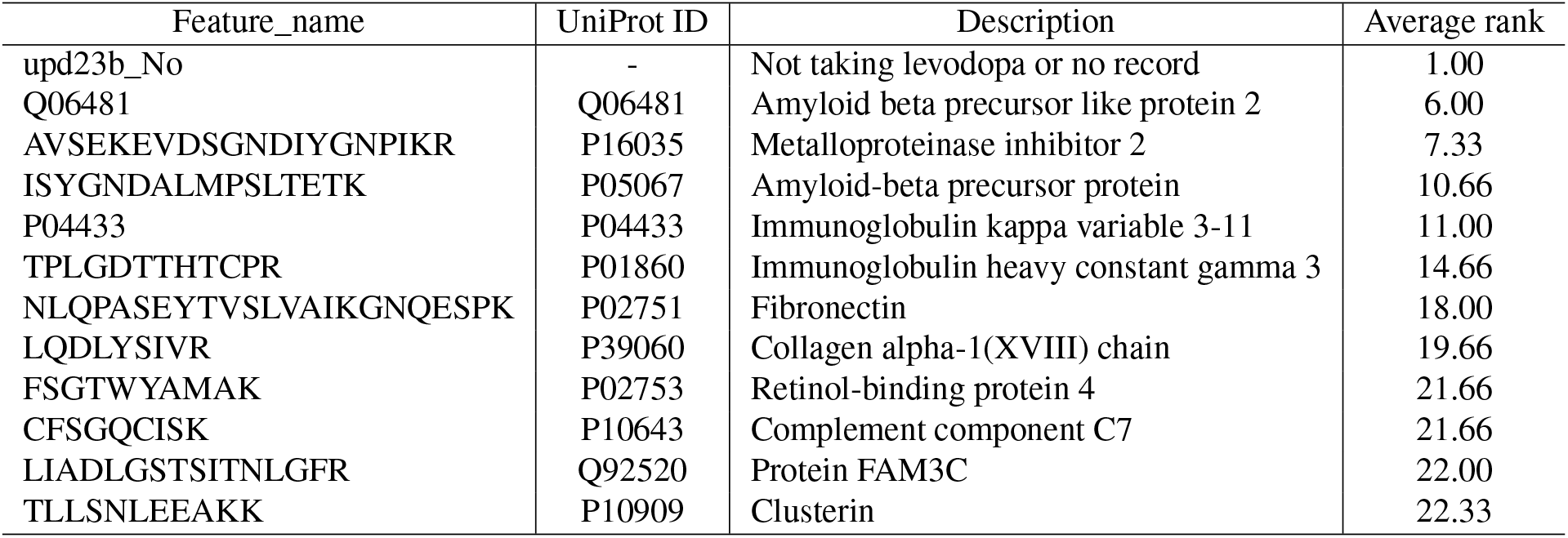
List of high-ranked importance features and their corresponding descriptions

It turns out that the usage of Levodopa has the most substantial impact on the results based on the rankings obtained through all interpretability methods. Furthermore, We have identified 11 proteins and peptides deemed important as a result. Figure 2 provides a 3-D visualization of some example proteins and peptides identified to be associated with Parkinson’s disease or neuronal degenerative diseases like Alzheimer’s in the UniProt dataset. Notably, since a peptide is part of a protein, it has been highlighted separately in red within the protein it is part of.

**Figure 1.**
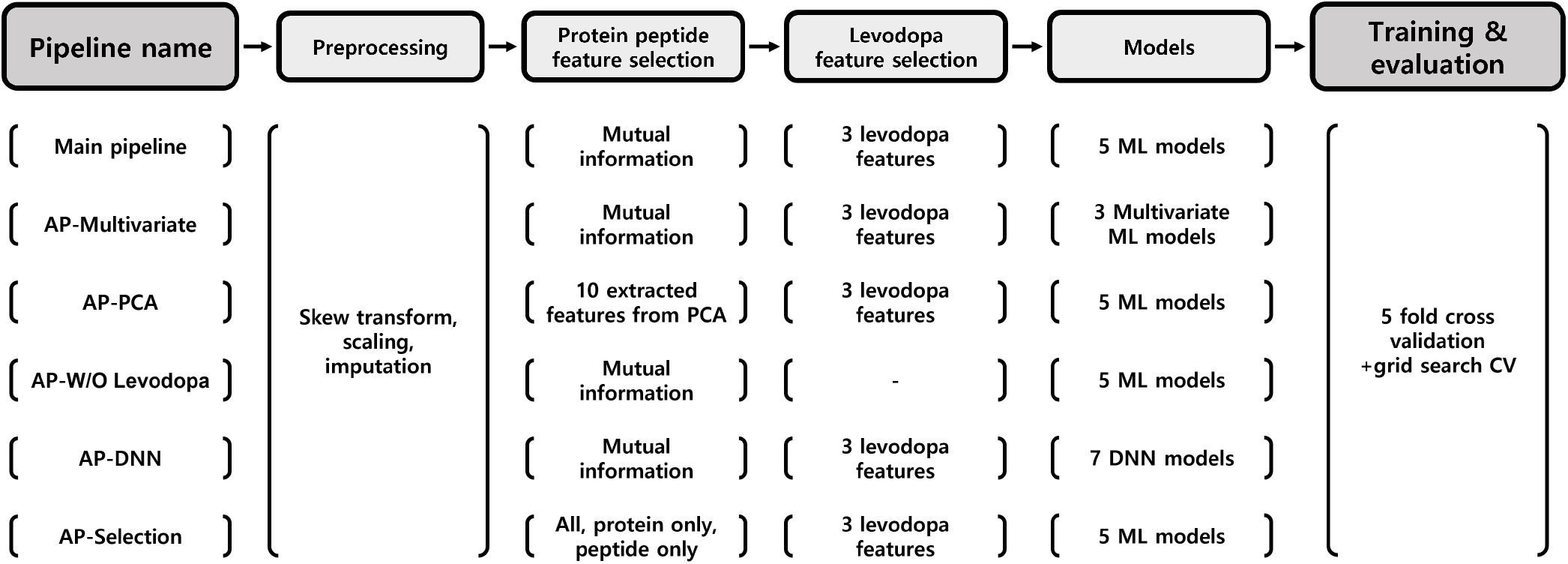
Overview of the processes of six pipelines including main and AP pipelines for performance comparison.

**Figure 2.**
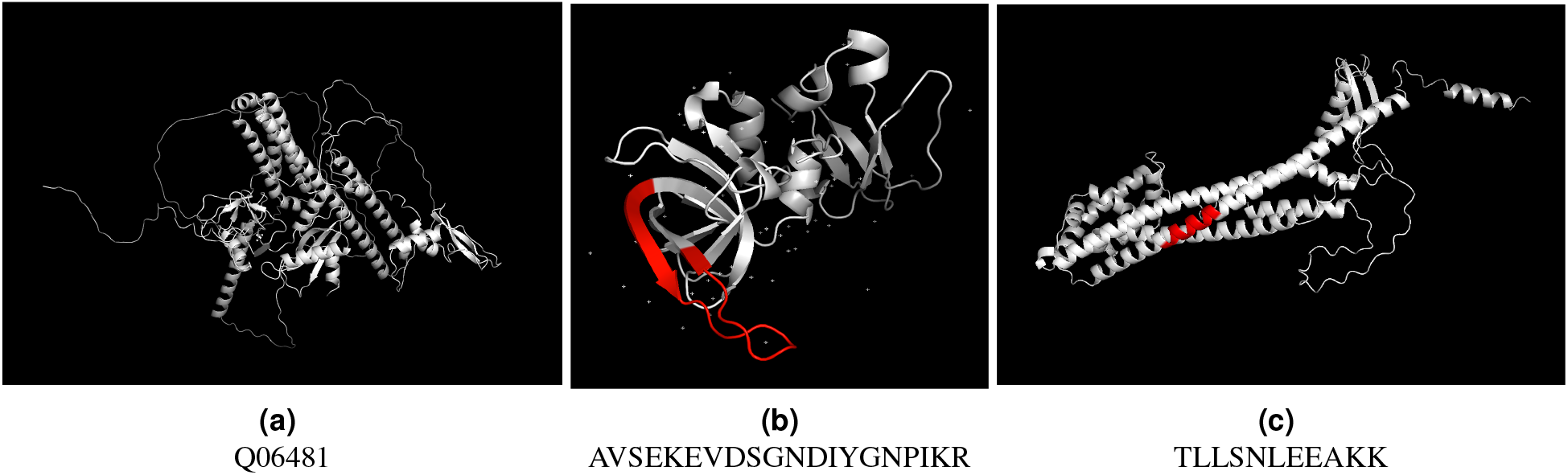
Example of 3-D structures of the top-ranked proteins and peptides. Peptides are indicated in red as part of their respective proteins.

### Comparing the Performance of Different Pipelines

Unlike in classification tasks, where the model’s performance can be intuitively gauged using accuracy or similar metrics, regression models are evaluated based on error metrics. This makes the assessment of the quality of results somewhat indirect and more challenging. To address this issue, we have implemented and analyzed five alternative pipelines, comparing their performance against our primary pipeline.

Figure 3d presents a comparison of the Average SMAPE between our primary pipeline and four additional pipelines (APs). The Average SMAPE offers a quantifiable measure for comparing the relative performance of the various pipelines. Our pipeline outperforms the other four, achieving the lowest Average SMAPE.

**Figure 3.**
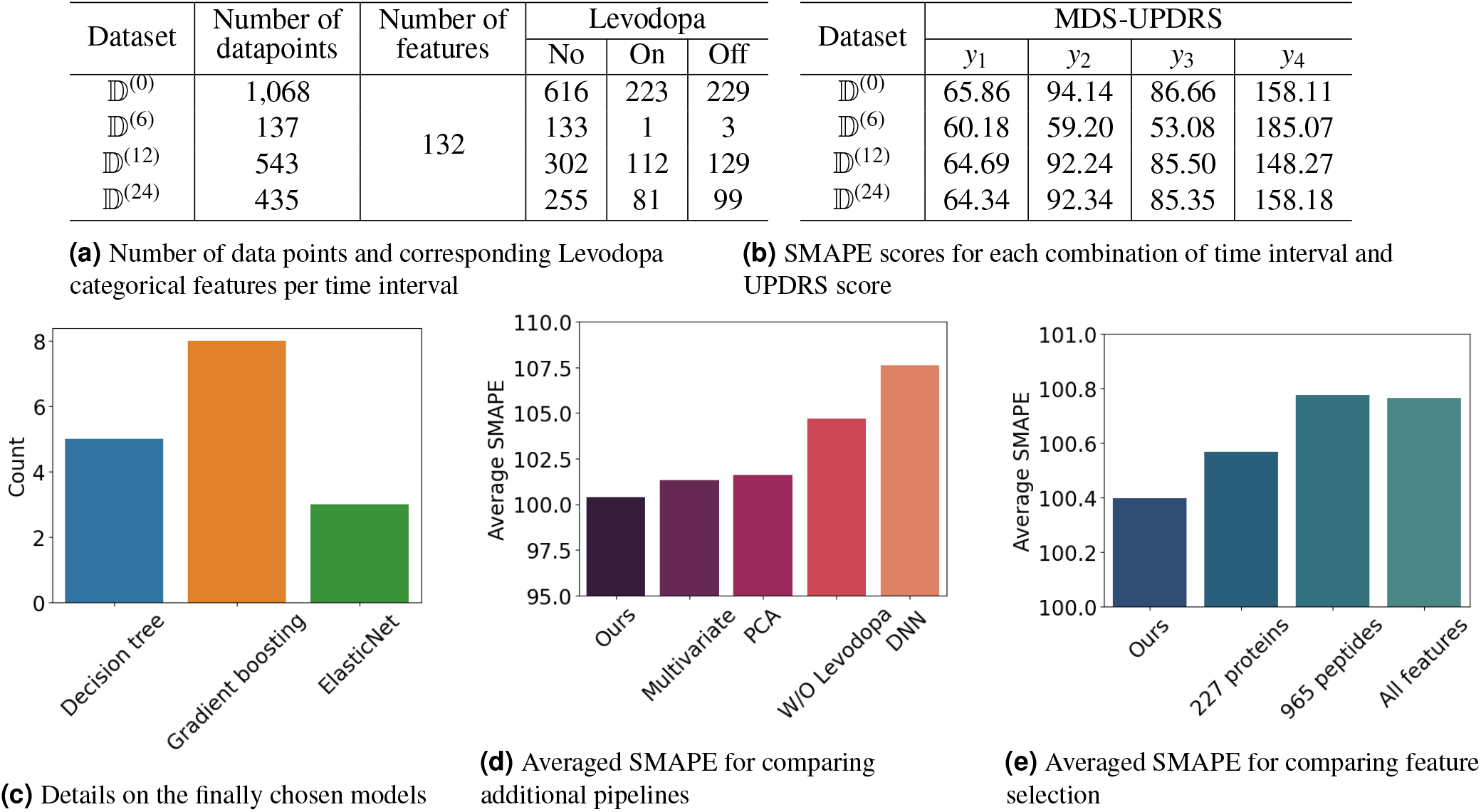
Overall results

Figure 3e illustrates the implications of excluding feature selection via Mutual Information (MI). We evaluated the relative Average SMAPE under three different conditions: when all proteins are utilized as input (*227 Proteins*), when all peptides are used as input (*965 Peptides*), and when all features are inputted (*All*). Our feature selection strategy consistently yields the lowest relative Average SMAPE, indicating superior predictive accuracy.

## Discussion

Our pipeline identified 2 proteins and 9 peptides as potential biomarkers for Parkinson’s disease, and found that Levodopa usage has the most significant impact on the results based on the average rankings obtained through all interpretability methods. Nine of the selected proteins and peptides were associated with neuronal degeneration and Alzheimer’s disease. All of the identified proteins and peptides were also associated with cancer. A summary of these associations can be found in the [Supplementary material (2)](https://docs.google.com/spreadsheets/d/1TAIsb41U-mO0Ot5ZYPxYIglemiErXkhodiW25JWI90E/edit?usp=sharing), while a more detailed description of each individual protein and peptide is provided in the [Supplementary material (3)](https://docs.google.com/document/d/1IKTu2hL0SFZJHU2AYKur6sSjRn5mKeJm/edit?usp=sharing&ouid=101633548124306174896&rtpof=true&sd=true). The 3D structures of the 129 MI-selected proteins and peptides, along with their explanations, can be conveniently explored using our structure viewer program. This program is accessible in [Supplementary material (5)](https://colab.research.google.com/drive/1EcqZmy0QL4QFJoIVa65RsttOQz9lxcg6?usp=sharing#scrollTo=sSNJplQ-iCw5).

Comparing our pipeline with four other APs revealed that our approach achieved the lowest Average SMAPE, indicating that the combination of MI-based feature selection and the selected regression models provided better performance in predicting UPDRS scores. Additionally, our feature selection strategy outperformed cases where all proteins, all peptides, or all features were used as input without selection. The code for all APs is available in the [Supplementary material (6)](https://drive.google.com/file/d/1wSegYfF3FrWR25WNYQTdqCoh6ySTiQLo/view?usp=share_link)

Our study offers several important contributions to the existing body of research on Parkinson’s disease. First, by focusing on a regression problem rather than a classification problem, our pipeline allows for prognostic prediction, which can be useful in tracking the progression of the disease and informing treatment decisions. Second, our MI-based feature selection approach efficiently reduces the dimensionality of the input data, improving model performance and interpretability. Finally, the combination of three interpretability methods increases the reliability of the identified candidate biomarker proteins and peptides.

Although our study makes interesting contributions, several limitations require further investigation. The first limitation is the small sample size, which raises concerns about the reliability of our results. While we had a sample size of 247 patients and 1,068 CSF-MS tests, it is important to note that we considered 1,195 features. Additionally, while the 𝔻^(6)^ had the lowest SMAPE, its reliability remains questionable. Another limitation is that even though our problem has the property of time difference, we did not use models for time series, such as LSTM. However, as confirmed in AP-ANN, deep learning-based models require a large amount of data to generalize.

To address these limitations, future research could focus on validating our pipeline, the identified biomarkers, and new time series-based prediction models in larger, independent cohorts such as the Parkinson Progression Marker Initiative [37] or Parkinson’s disease biomarkers program [38] datasets. These datasets contain long-term integrated treatment logs that could provide verification of the clinical utility of our pipeline. By testing our pipeline on these larger datasets, we can gain a better understanding of its generalizability and potential clinical impact. In addition, a multi-modal prediction approach will be possible by integrating other nucleotides, clinical, and blood data provided by those datasets.

In conclusion, our machine learning pipeline provides a promising approach for identifying candidate biomarker proteins and peptides in Parkinson patients using CSF-MS data. The proposed pipeline demonstrates the lowest average SMAPE and stable interpretability, offering valuable insights for understanding the progression of Parkinson’s disease and guiding clinical decision-making.

## Supporting information

Supplementary material (1) Machine learning configurations and results

Supplementary material (2) Summary of important Proteins and Peptides

Supplementary material (3) Detailed Description of Individual Proteins and Peptides

Supplementary material (4) Original Code of Our (main) Pipeline

Supplementary material (5) Visualization and Explanation Code for Proteins and Peptides

Supplementary material (6) Additional Pipeline Codes

## Acknowledgements

This research was supported by Ghent University Global Campus (GUGC) in Korea.The authors with like to thank the organizers of the AMP®-Parkinson’s Disease Progression Prediction competition and Professor Karsten Suhre of Weill Cornell Medicine - Qatar, who gently explained the mass spectrometry and dataset to us.

## Author contributions statement

HMP conceived and conducted the experiment(s), EK conducted the review of the manuscript, DM, MC, YK, and JI analyzed the results. AVM and WDN provided guidance and supervision. All authors reviewed the manuscript.

## Additional information

To include, in this order: **Accession codes** (where applicable); **Competing interests** (mandatory statement).

The corresponding author is responsible for submitting a [competing interests statement](http://www.nature.com/srep/policies/index.html#competing) on behalf of all authors of the paper.

This statement must be included in the submitted article file.

- **[Supplementary material (1)](https://docs.google.com/spreadsheets/d/1lu58P36kbP6XobipBoVMIDs0m656EJ0XLQcO2ZtNIi0/edit?usp=sharing): Machine learning configurations and results** - This supplementary material provides an overview of all machine learning pipelines used in the study along with the hyperparameters that were tested for each pipeline. Additionally, this material includes the detailed SMAPE results obtained by each pipeline.
- **[Supplementary material (2)](https://docs.google.com/spreadsheets/d/1TAIsb41U-mO0Ot5ZYPxYIglemiErXkhodiW25JWI90E/edit?usp=sharing): Summary of important Proteins and Peptides** - This supplementary material provides a summary of the 11 identified proteins and peptides and their associations with Parkinson’s disease and other neurodegenerative diseases, as well as cancer. In addition, the supplementary data contains a comprehensive ranking of the 132 features used in the model trainings.
- **[Supplementary material (3)](https://docs.google.com/document/d/1IKTu2hL0SFZJHU2AYKur6sSjRn5mKeJm/edit?usp=sharing&ouid=101633548124306174896&rtpof=true&sd=true): Detailed Description of Individual Proteins and Peptides** - This supplementary material provides a more detailed description of each of the 11 identified proteins and peptides, including their molecular characteristics, functional roles, and associated diseases.
- **[Supplementary material (4)](https://colab.research.google.com/drive/1zH3EqbLTioKL4G_TGM0X0U4tBKY1i3e5?usp=sharing): Original Code of Our (main) Pipeline** - This supplementary material contains the original code of our machine learning pipeline, implemented using Python and available through Google Colab. Users can run the code and adapt it for academic purposes. We have also provided pre-calculated inputs to save time and computational resources, allowing users to skip the initial data preprocessing steps if desired.
- **[Supplementary material (5)](https://colab.research.google.com/drive/1EcqZmy0QL4QFJoIVa65RsttOQz9lxcg6?usp=sharing#scrollTo=sSNJplQ-iCw5): Visualization and Explanation Code for Proteins and Peptides** - This supplementary material is a searchable code that displays the 3-D structure and explanations of all proteins referenced in the experiment.
- **[Supplementary material (6)](https://drive.google.com/file/d/1wSegYfF3FrWR25WNYQTdqCoh6ySTiQLo/view?usp=share_link): Additional Pipeline Codes** - This supplementary material includes the codes used in the experiment along with some comments.

