## Supplementary material (3) Detailed Description of Individual Proteins and Peptides for "Discovering Biomarker Proteins and Peptides for Parkinson’s Disease Prognosis Prediction with Machine Learning and Interpretability Methods"

**1. Q06481, Amyloid beta precursor-like protein 2 (APLP2)**

- Functions in neural development and synaptic transmission in the brain
- Thiamine-deficient brains show damage and accumulation of APP and APLP2 immunoreactivity
- Suggested that APP and APLP2 accumulation may be a compensatory response to neuronal damage
- Clusters of APP/APLP2 resemble Alzheimer's amyloid plaques
- Overexpression in cancer cells
- Dysregulation promotes tumor cell proliferation, migration, and invasion
- Abnormal processing leads to amyloid-beta peptide formation and Alzheimer's disease development

Amyloid precursor protein and amyloid precursor-like protein 2 in cancer - PMC (nih.gov)

Novel neuritic clusters with accumulations of amyloid precursor protein and amyloid precursor-like protein 2 immunoreactivity in brain regions damaged by thiamine deficiency. - PMC (nih.gov)

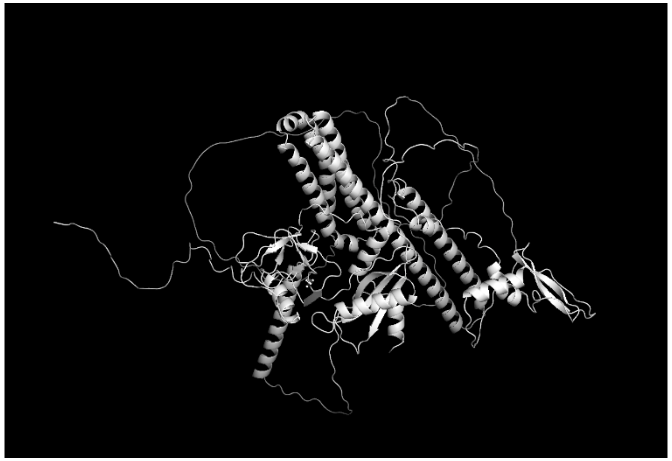

**2. P16035, Metalloproteinase inhibitor 2 (TIMP-2)**

- Belongs to the tissue inhibitors of metalloproteinases (TIMPs) family
- Inhibits matrix metalloproteinases (MMPs) activity, enzymes that break down proteins in the extracellular matrix (ECM)
- MMPs are involved in tissue remodeling, repair, and various pathological conditions (cancer and arthritis)
- Inhibits angiogenesis by affecting MMPs, which degrade ECM proteins and release angiogenic factors
- Potential therapeutic applications in cancer with aberrant angiogenesis
- TIMPs are up-regulated in the brain tissue of patients with Alzheimer's Disease

Increased plasma levels of matrix metalloproteinase-9 in patients with Alzheimer's disease - ScienceDirect

TIMP-2: an endogenous inhibitor of angiogenesis - ScienceDirect

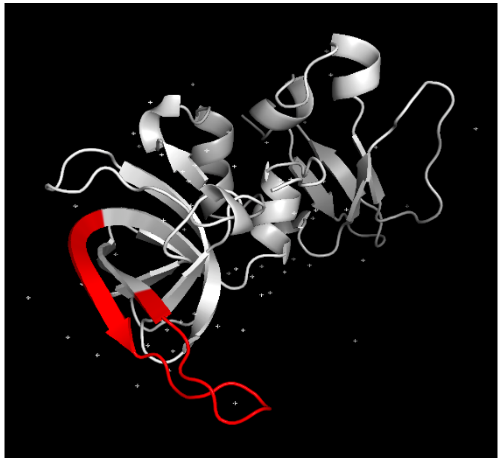

**3. P05067, Amyloid-beta precursor protein (APP)**

- Transmembrane proteins found in plasma membranes of various cells, including neurons
- Role in suppressing or reducing the synaptic transmission
- Linked to the accumulation of beta-amyloid plaques in the brain
- beta-amyloid peptide accumulation can form insoluble aggregates in the brain, which is considered a key factor in Alzheimer's disease pathogenesis
- Down-regulated in cancer cells

[Amyloid precursor protein and amyloid precursor-like protein 2 in cancer - PMC (nih.gov)]

[Amyloid-beta precursor protein processing in neurodegeneration - ScienceDirect]

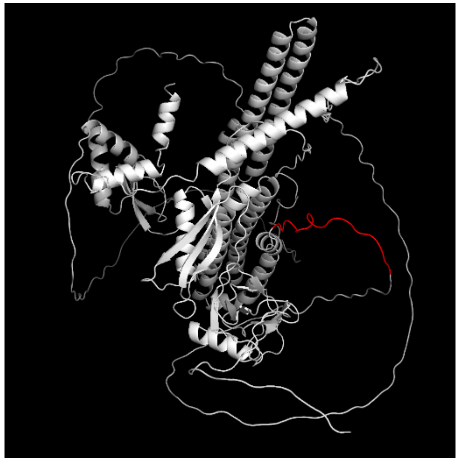

**4. P04433, Immunoglobulin kappa variable 3-11 (IGKV3-11)**

- Codes for a variable region of the kappa light chain in immunoglobulins (antibodies)
- Member of the human IGKV3 gene family
- Highly diverse and frequently used kappa light chain gene family in humans
- May play a role in the development of autoimmune diseases (e.g., rheumatoid arthritis)

A murine Ig light chain transgene reveals IGKV3 gene contributions to anti-collagen types IV and II specificities - PMC (nih.gov)

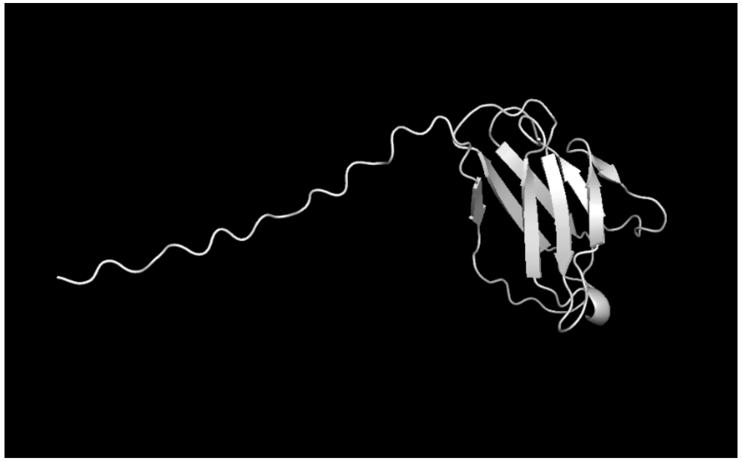

**5. P01860, Immunoglobulin heavy constant gamma 3 (IGHG3)**

- Encodes for the gamma 3 constant region of the heavy chain of immunoglobulin G (IgG) antibody
- The antibody generated by IGHG3 interact with beta-amyloid
- Associated with neuroinflammation in Alzheimer's disease
- IGHG3 level detected in systemic lupus erythematosus (SLE) which is an autoimmune disease
- Overexpressed IGHG3 in cases of pulmonary Yin deficiency syndrome (PYD), hyperactivity of fire due to Yin deficiency syndrome (HFYD), and deficiency of Qi and Yin syndrome (DQY).

Serum protein gamma-glutamyl hydrolase, Ig gamma-3 chain C region, and haptoglobin are associated with the syndromes of pulmonary tuberculosis in traditional Chinese medicine | SpringerLink

IJMS | Free Full-Text | Increased Immunoglobulin Gamma-3 Chain C in the Serum, Saliva, and Urine of Patients with Systemic Lupus Erythematosus (mdpi.com)

Whole exome sequencing study identifies novel rare and common Alzheimer's-Associated variants involved in immune response and transcriptional regulation | Molecular Psychiatry (nature.com)

- 

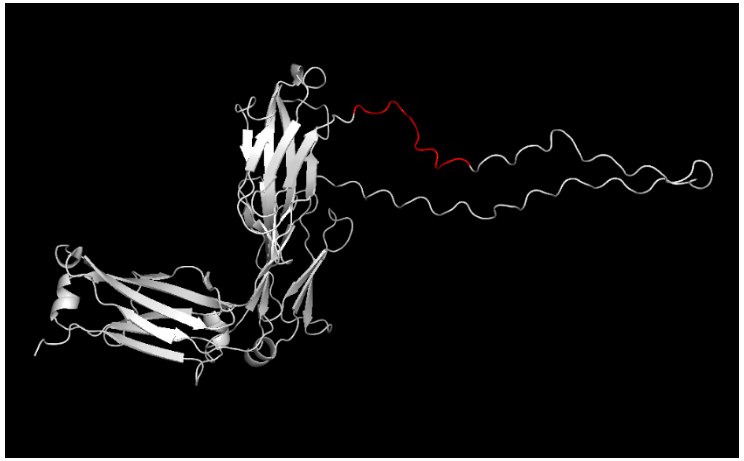

**6. P02751, Fibronectin (FN1)**

- Glycoproteins found in the extracellular matrix (ECM) of various tissues and organs
- Critical role in tissue development and repair
- Regulate angiogenesis in endothelial cells of the central nervous system
- Aberrant expression is associated with tumor growth, cancer, cardiovascular disease, and metastasis.
- Considered as a biomarker for assessing Alzheimer's disease risk

Targeting fibronectin for cancer imaging and therapy - Journal of Materials Chemistry B (RSC Publishing)

Molecular Status of Plasma Fibronectin as an Additional Biomarker for Assessment of Alzheimer's Dementia Risk | Dementia and Geriatric Cognitive Disorders | Karger Publishers

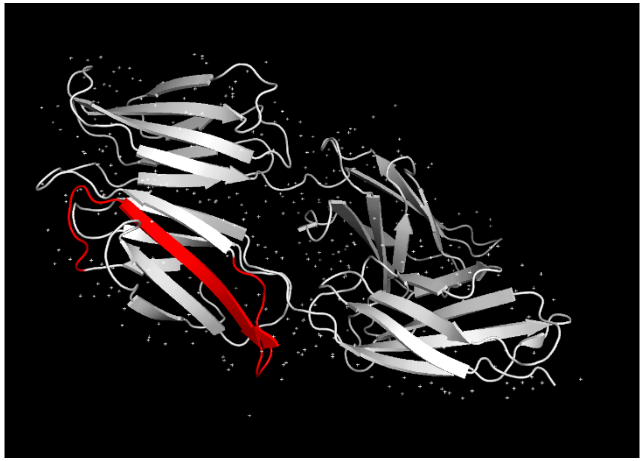

**7. P39060, Collagen alpha-1(XVIII) chain (COL18A1)**

- Encodes for a type of fibrous protein, collagen
- Plays a critical role in tissue structure, angiogenesis regulation and closure of the neural tube
- Mutations or dysregulation linked to several diseases, including Knobloch syndrome which is an autosomal recessive disorder and tumor growth

Altered expression of angiogenesis-related placental genes in pre-eclampsia associated with intrauterine growth restriction: Gynecological Endocrinology: Vol 23, No 6 (tandfonline.com)

Collagen XVIII, containing an endogenous inhibitor of angiogenesis and tumor growth, plays a critical role in the maintenance of retinal structure and in neural tube closure (Knobloch syndrome) | Human Molecular Genetics | Oxford Academic (oup.com)

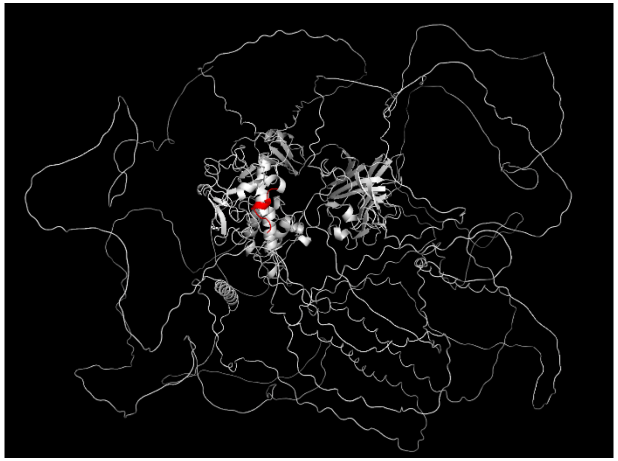

**8. P02753, Retinol-binding protein 4 (RBP4)**

- Primarily produced by the liver and circulates in blood
- Binds to and transports retinol (vitamin A) to other body tissues
- Involved in regulation of glucose and lipid metabolism
- Potentially plays a role in insulin resistance and impaired glucose tolerance
- Possible link between obesity and cancer
- Elevated RBP4 protein levels in the brain of mouse models with Alzheimer's disease

Is retinol binding protein 4 a link between adiposity and cancer? (degruyter.com)

Susceptibility to diet-induced obesity and glucose intolerance in the APP SWE/PSEN1 A246E mouse model of Alzheimer's disease is associated with increased brain levels of protein tyrosine phosphatase 1B (PTP1B) and retinol-binding protein 4 (RBP4), and basal phosphorylation of S6 ribosomal protein | SpringerLink

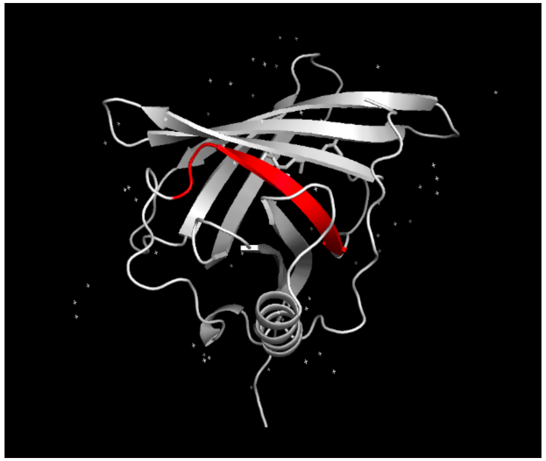

**9. P10643, Complement component C7 (C7)**

- Part of the complement system, critical for immune response to pathogens
- C7 acts as a potential tumor suppressor in relation to tumor progression and prognosis
- Complement activation products acting as an inflammatory mediator colocalize with beta-amyloid deposits in Alzheimer's disease (AD) brain
- Early complement components C1, C4, and C3 were found in neuritic plaque, and late complement components C7, C9, and membrane attack complex (MAC) were present in small quantities or undetected in Alzheimer's disease brain, which implies that the action of early complement components act as inflammatory mediator rather than late complement components or MAC in Alzheimer's disease.

The role of complement and activated microglia in the pathogenesis of Alzheimer's disease - ScienceDirect

Complement component 7 (C7), a potential tumor suppressor, is correlated with tumor progression and prognosis | Oncotarget

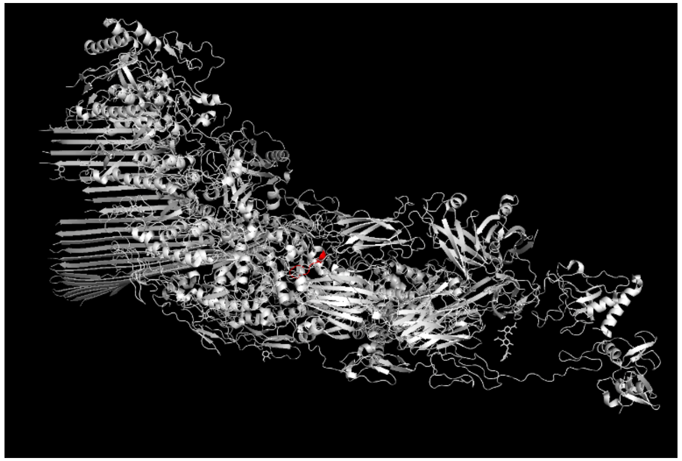

**10. Q92520, Protein FAM3C (FAM3C)**

- Member of FAM3 family
- Primarily expressed in the pancreas, regulates insulin secretion from pancreatic beta cells
- Role in reducing the production of beta-amyloid by affecting the β-secretase
- Involved in various physiological processes (inflammation, metastasis, and cell proliferation)
- Dysregulation associated with several diseases (obesity, type 2 diabetes, breast cancer, and colon cancer)
- Decrease of FAM3C expression in the brains of Alzheimer's disease patients

FAM3C: an emerging biomarker and potential therapeutic target for cancer | Biomarkers in Medicine (futuremedicine.com)

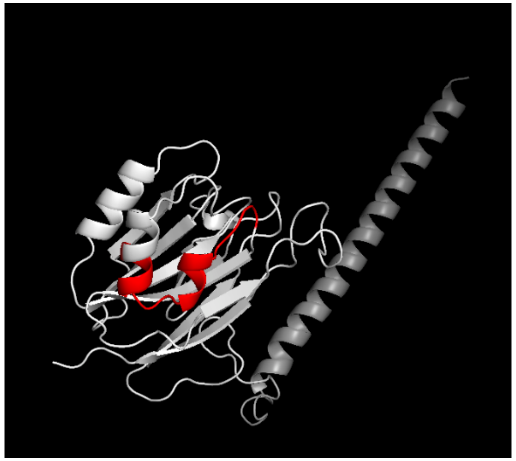

**11. P10909, Clusterin (CLU)**

- Clusterin binds to megalin receptors and facilitates endocytosis within glial cells to remove amyloid-β peptides and fibrils
- Dysregulation of expression associated with various diseases (Alzheimer's disease, cancer, kidney disease)
- In cancer, implicated in the resistance of cancer cells to chemotherapy and radiation therapy
- Increased clusterin levels found in brain and cerebrospinal fluid of Alzheimer's disease patients

Clusterin: A forgotten player in Alzheimer's disease - ScienceDirect

Clusterin inhibition mediates sensitivity to chemotherapy and rad...: Ingenta Connect

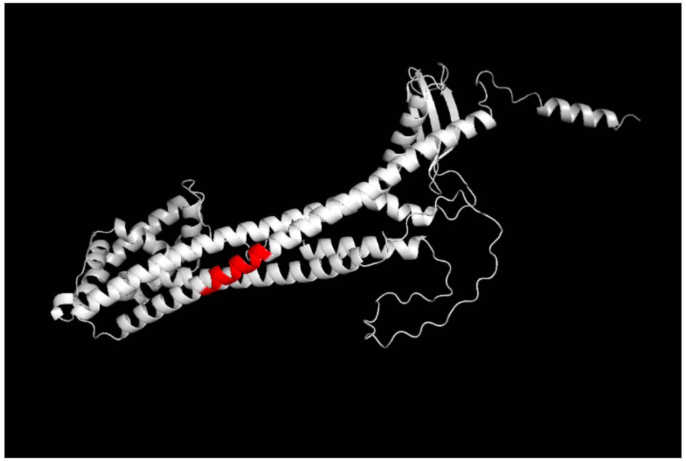
