## Supplementary material (4) Original Code of Our (main) Pipeline for "Discovering Biomarker Proteins and Peptides for Parkinson’s Disease Prognosis Prediction with Machine Learning and Interpretability Methods": Supplementary material (4) Original Code of Our (main) Pipeline.ipynb

```
# %% [markdown]
# Discovering Biomarker Proteins and Peptides for Parkinson’s Disease Prognosis Prediction with Machine Learning and Interpretability Methods

The purpose of this code is to provide a detailed explanation of the methodology presented in the research paper of the same name and share an example code for interested individuals. Specifically, this code demonstrates the process of identifying 11 candidate biomarkers from the AMP®-Parkinson's Disease Progression Prediction dataset. 

To enhance usability, we have made the intermediate results generated during the code execution downloadable. This allows users to skip the intermediate steps if needed and directly proceed to the desired part of the code. 

While we have taken utmost care to ensure code reproducibility by adjusting the random seed, it is important to note that code execution may encounter challenges due to version differences. If you encounter any difficulties, please do not hesitate to contact us at for further assistance.

## Introduction

Parkinson's Disease is a progressive disorder of the nervous system that affects movement. There is currently no cure for Parkinson's, but early diagnosis and treatment can help manage symptoms and improve quality of life. In this Kaggle competition, we are tasked with predicting the progression of Parkinson's disease using a dataset of patient clinical and demographic information.


## Table of Contents

1. Understanding dataset (EDA)
- 1.1. X: Protein abundance values derived from cerebrospinal fluid (CSF) mass
- 1.2. Y: Movement Disorder Society Unified Parkinson's Disease Rating Scale (MDS-UPDRS)

2. Pre-processing
- 2.1. Downloading and importing packages
- 2.2. Read train csvs
- 2.3. Mergning dataframes
- 2.4. EDA using spreadsheet and raw eye
- 2.5. Skewness checking and removing
- 2.6. Calculating scalers
- 2.7. Checking type of missing data and imputing
- 2.7.1. Imputation of feature (X) values
- 2.7.2. Imputation of target (Y) values

3. Feature selection, feature engineering and Dataset dividing
- 3.1. Feature selection using mutual information 
- 3.2. Onehot encoding `upd23b_clinical_state_on_medication` and mergning all the data 
- 3.3. Dataset dividing for time series

4. Training and evaluation 
- 4.1. Dataset checking before training
- 4.2. Training models with hyperparamter cross validation
- 4.3. Reading results
- 4.4 Setting final models

5. Finding candidate biomarkers
- 5.1. Calculating three different interpretable AI methods
- 5.2 Summarizing the all results


We will begin by exploring the dataset and performing data cleaning and preprocessing. Then, we will engage in feature engineering to extract relevant information from the dataset and select appropriate features for modeling. Next, we will develop and evaluate various machine learning models, selecting the most appropriate one for deployment. Finally, we will generate predictions using our chosen model and interpret the results. We will conclude by summarizing our findings and discussing potential areas for future work.


# %% [markdown]
## 1. Understanding dataset (EDA)


This chapter aims to identify the problem definition in the AMP-Parkinson's Disease Progress competition and understand the characteristics of the dataset accordingly.

The competition organizers are interested in **predicting** the Unified Parkinson's Disease Rating Scale (UPDRS) score, which consists of **4 real values (multivariate regression)**, using around **1,200 types of protein/peptide abundance inputs**. Additionally, the problem has a **time series nature**, meaning that participants are required to predict the patient's UPDRS status at the current ($t_0$), 6+month ($t_6$), 12+month ($t_{12}$), and 24+month ($t_{24}$) time points using the patient's mass spectrometry input. It is not yet clear whether additional protein/peptide abundance values are given after $t_0$, the point at which the input begins.

# %% [markdown]
### 1.1. X: Protein abundance values derived from cerebrospinal fluid (CSF) mass spectrometry

The input to this data is **protein abundance values** derived from **mass spectrometry** readings of **cerebrospinal fluid (CSF)** samples gathered from several hundred patients. In mass spectrometry, the sample is ionized and the resulting ions are separated based on their mass-to-charge ratio using electric and/or magnetic fields. The separated ions are then detected and their abundance is measured, allowing for the identification and quantification of the molecules in the sample. In the case  of CSF samples, mass spectrometry can be used to **identify and quantify the various proteins present in the fluid**, which can provide valuable information about the health status of the patients. By comparing the protein abundance values across different patients, researchers can **identify potential biomarkers of disease or other conditions**, which can ultimately help in the development of  new diagnostic and treatment strategies.
 
 
 

# %% [markdown]
### 1.2. Y: Movement Disorder Society Unified Parkinson's Disease Rating Scale (MDS-UPDRS)

**MDS-UPDRS**, or Movement Disorder Society Unified Parkinson's Disease Rating Scale, is a widely used tool in clinical practice and research for **evaluating the severity and progression of Parkinson's Disease**. This neurodegenerative disorder primarily affects movement, and MDS-UPDRS assesses both motor and non-motor symptoms and monitors changes over time.

#### MDS-UPDRS Parts and Assessment

MDS-UPDRS is divided into several parts, each of which assesses a different aspect of Parkinson's Disease:

-   **Part I - Non-Motor Aspects of Experiences of Daily Living:** This section evaluates non-motor aspects of Parkinson's Disease, such as mood, cognition, and sleep.
-   **Part II - Motor Aspects of Experiences of Daily Living:** This section assesses the impact of Parkinson's Disease on daily activities, such as dressing, hygiene, and eating.
-   **Part III - Motor Examination:** This section assesses the severity of Parkinson's Disease motor symptoms, such as tremors, rigidity, and bradykinesia.
-   **Part IV - Motor Complications:** This section evaluates the presence and severity of motor complications related to Parkinson's Disease treatment, such as dyskinesias and motor fluctuations.

From the part I - III, there are several items that are rated on a scale of 0 to 4 points. The higher the score, the higher the likelihood of Parkinson's disease. The sum of these points is used to determine the overall score for that part, with a maximum possible score of 52 points and a minimum score of 0 points. 

#### MDS-UPDRS Administration

MDS-UPDRS is administered by a trained healthcare professional and typically **takes 30-60 minutes** to complete. The results of the assessment are then used to guide treatment decisions and monitor disease progression over time.


# %% [markdown]
## 2. Pre-processing


# %% [markdown]
### 2.1. Downloading and importing packages

# %% [code]
!pip install fancyimpute
!pip install gdown


# %% [code]
import pandas as pd
import seaborn as sns
import os, code 
import numpy as np
from sklearn.preprocessing import StandardScaler
import pickle
import scipy.stats as stats
import matplotlib.pyplot as plt
import pprint
import copy, random
from scipy.stats import boxcox,anderson, shapiro
from fancyimpute import SoftImpute


# Set random seed for reproducibility
seed = 42
random.seed(seed)
np.random.seed(seed)


# %% [markdown]
### 2.2. Read train csvs
There are **248 unique patient IDs**, **227 unique Uniprot IDs**, and **968 unique peptide IDs**.

In other words, since there are more than five times more available protein & peptide features (227 + 968) than the amount of usable data (227), we need to be serious about feature selection.


# %% [code]
df_clinical = pd.read_csv("/kaggle/input/amp-parkinsons-disease-progression-prediction/train_clinical_data.csv")
df_protein = pd.read_csv("/kaggle/input/amp-parkinsons-disease-progression-prediction/train_proteins.csv")
df_pebtide = pd.read_csv("/kaggle/input/amp-parkinsons-disease-progression-prediction/train_peptides.csv")

petient_list = list(df_clinical["patient_id"].unique())
uniprot_list = list(df_protein["UniProt"].unique())
peptide_list = list(df_pebtide["Peptide"].unique())

print("# of unique patient Ids:", len(petient_list))
print("# of unique Uniprot (protein) Ids:", len(uniprot_list))
print("# of unique Peptide Ids:", len(peptide_list))


# %% [markdown]
### 2.3. Mergning dataframes


Our goal is to find out the correlation between UPDRS scores and proteins & peptides and to create artificial intelligence that can learn it. However the data is spread across three files, it is challenging to use and observe intuitively. Therefore, in this code, we will merge these three files (or dataframes) into a single dataframe and save the integrated dataset as **"integrated_dataset.csv"** locally. This may take some time, so you can download the data through the download code after the main code


# %% [code]
# This takes quite a long time. It would be better to download the processed dataset.
"""
total_list=  list(df_clinical.columns) + uniprot_list + peptide_list
df = pd.DataFrame(columns=total_list)

for idx, visit_id in enumerate(df_clinical['visit_id']):
	if idx % 100 == 0:
		 print(idx,"/",len(df_clinical))
	df_cli = df_clinical[df_clinical['visit_id']==visit_id]
	df_pro = df_protein[df_protein['visit_id'] == visit_id]
	df_pep = df_pebtide[df_pebtide['visit_id'] == visit_id]
	dic_protein = {}
	pro_dic = {key:value for key, value in zip(df_pro['UniProt'], df_pro['NPX'])}
	pep_dic = {key:value for key, value in zip(df_pep['Peptide'], df_pep['PeptideAbundance'])}

	pro_dic_out = {item : (pro_dic[item] if item in pro_dic else np.nan)  for item in uniprot_list}
	pep_dic_out = {item : (pep_dic[item] if item in pep_dic else np.nan)  for item in peptide_list}

	df_cli = df_cli.iloc[0].to_dict()
	df_cli.update(pro_dic_out)
	df_cli.update(pep_dic_out)
	new_df = pd.DataFrame(df_cli, index=[0])
	df = pd.concat([df, new_df], ignore_index=True)
df.to_csv("integrated_dataset.csv", index=False)
"""

# %% [code]
#Downloading integrated_dataset.csv
!gdown --id "1Hz0sMjz2_xtw8JwdV0yb4r_33U0WRyAe"

#url = "https://drive.google.com/file/d/1Hz0sMjz2_xtw8JwdV0yb4r_33U0WRyAe/view?usp=sharing" 

# %% [markdown]
### 2.4. EDA using spreadsheet and raw eye

Let's do the most basic EDA. First of all, download the created `integrated_dataset.csv` and open the file using a spreadsheet of your choice (Excel, Google Spreadsheet, etc.).

If you check the X value, it appears that **CSF mass spectrometry was performed once every 6 months** between 0 and 48 months. Therefore, it can be seen that all numbers in the Protein and Peptide columns are NaN (Not a number) in the middle rounds, such as 55_3 and 55_9. In summary, CSF mass spectrometry has been performed on 1,068 rows, and upon viewing the dataframe, **substantially the dataframe contains 1068 rows × 1204 columns**. Also, **CSF mass spectrometry cannot detect all proteins and peptides.** For example, one of the peptides, `PALEDLR`, is also NaN at 55_0, but it can be confirmed that it is detected at 55_6 and 55_12.

If you check the Y value, you can see that there are a lot of missing values in `updrs_4`. This score is associated with dyskinesias and dystonia, which check impulsive muscle movements independent of the patient's will. What is unusual is the `upd23b_clinical_state_on_medication` column. This is a space to record the prognosis after applying of levodopa. **Levodopa** is converted to dopamine in the brain and increases the concentration of dopamine in the brain. It has been shown to be most effective for Parkinson's disease and particularly effective for stiffness and slow movements, but is known to have side effects. At this time, the column to be careful since **both ON/OFF means levodopa treatment has occurred**, and blank means no treatment. If there seems to be some positive effect after administration, it is marked as ON, otherwise it is marked as OFF, In the case of patient 55, levodopa was applied from the 9th month, and it seems to see positive effects until the 12th month.

Ahother interesting finding is that there is a lot of **difference in the values range of each column**. For example, protein `O00391` has values in the 10K range, but peptide `QKVEPLRAELQEGAR` has values in the 2.5M range.

Finally, we will **remove all the rows** for which this **CSF mass spectrometry has not been performed** and save them to the dataframe **'df_with_CSF'**. This is to remove unnecessary rows for which no checks have been performed.


# %% [code]
df = pd.read_csv("integrated_dataset.csv",index_col=False)

num_of_null = df[uniprot_list].isnull().sum(axis = 1)
print("Total number of rows in the integrated dataset", len(df))
print("# of CSF mass spectrometry was performed", len(df) - len(num_of_null[num_of_null == 227]))

display(df)


#Removing the row that does not have mass spectrometry data
num_of_null = df[uniprot_list].isnull().sum(axis = 1)
df_with_CSF = df[num_of_null != 227] 


# %% [markdown]
### 2.5. Skewness checking and removing

Skewness quantifies the asymmetry of a probability distribution. In a normal distribution, the mean, median, and mode are all equivalent, and the distribution is symmetric, resulting in a skewness of zero. Positive skewness signifies a longer tail on the positive side of the distribution compared to the negative side, while negative skewness indicates the opposite.

In machine learning, skewness can influence the performance of models, particularly when employing algorithms that assume a normal or symmetrical distribution of data. For instance, linear regression assumes the target variable follows a normal distribution, and any skewness present in the target variable may impact the model's effectiveness.

Consequently, **transforming the data to reduce skewness might enhance model performance**. In this case, we will employ **log, Box-Cox, and square root transformations** to mitigate the skewness of the dataset.


# %% [code]
feature_list = uniprot_list + peptide_list
df_features = df_with_CSF[feature_list]
transformations = {}

#1. Find skewed feature and applying log transformation
skewness = df_features.skew().sort_values(ascending=False)
skewed_features = skewness[skewness.abs() > 1].index.tolist()
print("# of skewed features / total features:\t",len(skewed_features), "/",len(feature_list))
print("skewed:", skewed_features)

df_features[skewed_features] = np.log1p(df_features[skewed_features])
transformations["1.log"] = copy.deepcopy(skewed_features)
skewness = df_features.skew().sort_values(ascending=False)
skewed_features = skewness[skewness.abs() > 1].index.tolist()
print("After log tranformations:\t\t",len(skewed_features), "/",len(feature_list))
print("skewed after log tranformation:", skewed_features)


#2. Apply Box-Cox transformation to further reduce the skewness for feature in skewed_features:

for feature_name in skewed_features:
    df_features.loc[:,feature_name], _ = boxcox(df_features[feature_name]+1)
transformations["2.boxcox"] = copy.deepcopy(skewed_features)
skewness = df_features.skew().sort_values(ascending=False)
skewed_features = skewness[skewness.abs() > 1].index.tolist()
print("After boxcox tranformations:\t\t",len(skewed_features), "/",len(feature_list))
print("skewed after box-cox tranformation:", skewed_features)

#3. Apply sqrt transformation to final reducing
df_features[skewed_features] = np.sqrt(df_features[skewed_features])
transformations["3.sqrt"] = copy.deepcopy(skewed_features)
skewness = df_features.skew().sort_values(ascending=False)
skewed_features = skewness[skewness.abs() > 1].index.tolist()
print("After square root tranformations:\t",len(skewed_features), "/",len(feature_list))
print("skewed after sqrt tranformation:", skewed_features)


#4 remove the columns that still has sknewness
df_features = df_features.drop(columns=skewed_features, axis=1)
feature_list = df_features.columns.tolist() #update the feature list
transformations["4.remove"] = skewed_features


#5. Save those feature transformations info for testset
pickle.dump(transformations,open("transformations.pkl", "wb"))


# %% [markdown]
### 2.6. Calculating scalers


Let's look at "df_features" with skewness correction completed. It is evident that the **feature scale difference has become significantly large**. A tranformation, such as a log tranformation, exponentially changes the size of a value (ex, log(10000) = 4). For example, the value for protein `O00533` has a mean of 509,201 and a standard deviation of 233,573. Conversely, `O14791` with tranformation applied has an average of 7.8 and a standard deviation of only 0.4. 

In data analysis, machine learning, and statistical modeling, different scales of input values **can lead to several problems**, such as some features dominating the final predictions. 

To address these problems, we would **apply standard scaling to the input features** so that they have a consistent scale. Normalization or standardization can help ensure that all features are given equal importance and that the model is not biased by the scale of the input values. 

Standardization is a technique used to transform data so that it has a **mean of zero and a standard deviation of one**. This is achieved by subtracting the mean from each data point and then dividing by the standard deviation.


# %% [code]
#NaN checking through raw eye and spreadsheet
df_features = df_features.drop(df_features.filter(regex='Unnamed').columns, axis=1)
print("# of NaN:",df_features.isna().sum().sum())
display(df_features)

print(df_features['O00533'].describe())
print("---------------------")
print(df_features['O14791'].describe())

# %% [code]
#Calculating scalers and save csv and pickle files

scalers = {item: StandardScaler() for item in uniprot_list}

for col_name in df_features.columns:
	scalers[col_name] = StandardScaler()
	df_features.loc[:,col_name] = scalers[col_name].fit_transform(np.asarray(df_features[col_name]).reshape(-1, 1)).flatten()

pickle.dump(scalers, open("scalers.pkl", "wb"))
df_features.to_csv("df_features_scaled.csv", index=False)


# %% [markdown]
### 2.7. Checking type of missing data and imputing


In general, the types of missing values can be divided into the following three types: Missing completely at random (MCAR), Missing at random (MAR), and Missing not at random (MNAR).

The **missing value type** is determined by figuring out what kind of correlation the column including the missing value has with other columns. For example, let's say we want to predict a person's obesity (y) using height (X0), weight (X1) and sex (X(2)). If a large number of missing values occur in height, we can **divide all data points into `subset A` with all missing values and `subset B` without**, and then check whether other features are affected by this.

In other words, The type of missing value is possible to **find out whether there is a significant difference in weight, set and obesity between `subset A` and `subset B`** through a t-test (in the case of continuous variable), and a chi square test (in the case of categorical variable). If `subsets A` and `B` do not have significant differences in any feature, it is Missing completely at random (MCAR). If `subjects A and B` show a significant difference in obesity, it is Missing not at random (MNAR), and if there is a significant difference in weight or gender, which are other features other than obesity (y), it is Missing at random (MAR).

This information can be summarized in the table below:


| Missing Value Type | Definition | Characteristics |
| :---: | :---: | :---: |
| MCAR | Missing Completely at Random | No relationship between the missingness and other variables (X and y) |
| MAR | Missing at Random | Relationship between the missingness and other variables (X), but not the variable itself (y) |
| MNAR | Missing not at Random | Relationship between the missingness and the variable itself (y) or some unobserved factor related to the variable |


**Simple imputation (mean) is applied in** cases where missing values are Missing Completely At Random (**MCAR**) since this type of missingness is independent of any other feature dimension. However, Missing At Random (**MAR**) **and** Missing Not At Random (**MNAR**) missingness are related to the other features or target variables, and using simple imputation methods may cause correlation changes between features. 

# %% [markdown]
### 2.7.1. Imputation of feature (X) values

A total of 1192 features were identified, with an **average of 93 NaNs and a median of 10 NaNs per feature.** Additionally, **only 35 out of the 1192 features had zero NaNs**, indicating that there are **too many missing values** in this dataset **to remove by simply deleting data points**. Therefore, alternative methods must be considered for handling the missing values.

In this chapter, we will begin by **classifying the type of missing value** and imputing MCAR and MAR+MNAR separately. For **MCAR**, we will use **mean imputation**, while for imputing **MAR and MNAR**, we will utilize **SoftImpute** from the fancyimpute library.


# %% [code]
nan_count = df_features.isna().sum()
sns.histplot(nan_count)
print(nan_count.describe())  
print("Columns does not have NaN:",len(nan_count[nan_count ==0]))
plt.title("Number of NaN in each columns(protein and peptide)")
plt.show()

# %% [code]
#Checking whether each column (X) is related to other columns (X + Y)

import warnings
warnings.filterwarnings('ignore')

#removing the row that does not have mass spectrometry data
num_of_null = df[uniprot_list].isnull().sum(axis = 1)
df_with_CSF = df[num_of_null != 227] 

df_features = df_features[feature_list]
missing_counts = df_features.isnull().sum()
test_dictX = {}
test_dicty = {}
df_targets = df_with_CSF[["updrs_1","updrs_2","updrs_3","updrs_4"]]

for col in df_features.columns:

	missing = df_features[col].isnull()
	observed = ~missing
    #If the number of missing value is less than 30, treat as MCAR because it does not meet the minimum quantity of t test
	if sum(missing) <30: 
		test_dictX[col] = 0
		test_dicty[col] = 0
	else:
		# check for differences in other variables (X)
		other_cols = [c for c in df_features.columns if c != col]

		ret_list = []
		tar_list = []
		for other_col in other_cols:
			A = df_features[other_col][missing].dropna()
			B = df_features[other_col][observed].dropna()
			ret_list.append(stats.ttest_ind(A, B).pvalue < 0.05)

		test_dictX[col] = sum(ret_list)

        # check for differences in other variables (Y)
		for target in df_targets.columns:
			A = df_targets[target][missing].dropna()
			B = df_targets[target][observed].dropna()
			tar_list.append(stats.ttest_ind(A, B).pvalue < 0.05)
		test_dicty[col] = sum(tar_list)

		#print(col, sum(ret_list), sum(tar_list),sum(missing), sum(observed))
my_dic = {"test_dictX":test_dictX,"test_dicty":test_dicty}
with open('missing_values.pickle', 'wb') as file:
    pickle.dump(my_dic, file)


# %% [code]
#Downloading missing_values.pickle for fast run
#!gdown --id "1FAy7zCBTC-xHfgSjIzQMqKiPEzSPpJ3D"


# %% [code]
#Classifing the type of missing values in each column
df_targets = df_with_CSF[["updrs_1","updrs_2","updrs_3","updrs_4"]]

with open('missing_values.pickle', 'rb') as file:
    my_dic = pickle.load(file)    

test_dictX, test_dicty = my_dic["test_dictX"],my_dic["test_dicty"]

#fillcing MCAR and checking MAR and MNAR
filled_columns = []
MAR_columns = []
MNAR_columns = []
for col in df_features:
	if test_dictX[col] == 0 and test_dicty[col] == 0:
		#MCAR
		df_features[col].fillna(df_features[col].mean(), inplace=True)
		filled_columns.append(col)
	elif  test_dicty[col] == 0:
		MAR_columns.append(col)
	else:
		MNAR_columns.append(col)

mice =  SoftImpute(verbose=False)

df_combined = pd.concat([df_features, df_targets], axis=1)
print("# of MCAR:\t",len(filled_columns))
print("# of MAR:\t",len(MAR_columns))
print("# of MNAR:\t",len(MNAR_columns))

#impute MAR and MNAR missing valuee using softimpute
cols_to_impute = filled_columns+MAR_columns+MNAR_columns
imputed = mice.fit_transform(df_features[cols_to_impute])
df_features.loc[:,cols_to_impute] = imputed[:,:]
df_features.to_csv("df_features_scaled_imputed.csv", index=False)	

# %% [code]
#Downloading df_features_scaled_imputed.csv for fast run
#!gdown --id "1U_T_QWLeGzoXsZCMy9dChn-lZQFFSmPx"


# %% [markdown]
### 2.8.2. Imputation of target (Y) values

After examining the missing values in y, it was discovered that `updrs_3` had 10 missing values and updrs_4 had 499 missing values. The distribution of each updrs (excluding missing values) is displayed in the following figure using a KDE plot. **`updrs_4` has an average value close to 0 and a significant number of zeros**, hence NaN values were replaced with 0s. On the other hand, **`updrs_3`** has a very high maximum value and an average of 17, but since the number of **missing values is less than 1%**, which makes hard to test the type of missing values. Therefore we **replaced NaN values to 0** since the pick of the `updrs_3` is close to zero.

After studying the MDS-UPDRS, I found that **item 4 of the UPDRS questionnaire includes questions related to dyskinesias**, which are a known side effect of long-term levodopa treatment. Therefore, it is possible that item 4 only applies to patients taking levodopa, who have the upd23b_clinical_state_on_medication variable marked as ON/OFF. As a result, there may be **samll case that meets taking levodopa and having dyskinesias symtoms together**. Therefore I  **imputed NaN values in the updrs_4 column to 0**

# %% [code]
#removing the row that does not have mass spectrometry data
num_of_null = df[uniprot_list].isnull().sum(axis = 1)
df_with_CSF = df[num_of_null != 227] 

print('Missing values of updrs_1',sum(df_with_CSF['updrs_1'].isna()))
print('Missing values of updrs_2',sum(df_with_CSF['updrs_2'].isna()))
print('Missing values of updrs_3',sum(df_with_CSF['updrs_3'].isna()))
print('Missing values of updrs_4',sum(df_with_CSF['updrs_4'].isna()))


print(df_with_CSF[['updrs_1','updrs_2','updrs_3','updrs_4']].describe())

plt.subplots(figsize=(16,5),constrained_layout=True)
for i in range(1,5):
	plt.subplot(1,4,i)
	sns.kdeplot(df_with_CSF['updrs_'+str(i)], fill= True)
	new = df_with_CSF['updrs_'+str(i)]
	new = np.nan_to_num(new, nan=0)
	new = pd.Series(new.flatten(), index = df_with_CSF['updrs_'+str(i)].index)
	df_with_CSF.loc[:,'updrs_'+str(i)] = new	
 

# %% [markdown]
## 3. Feature selection, feature engineering and Dataset dividing


In Chapter 2, we worked on creating a comprehensive dataset by merging three CSV files provided in the competition, correcting skewness, scaling the data, and performing imputation to handle missing values. In this chapter, we will delve deeper into advanced data processing techniques such as feature selection, feature engineering, and dataset splitting for predicting time differences. These techniques will enable us to extract meaningful insights from the data and improve the accuracy of our predictions.


# %% [markdown]
### 3.1. Feature selection using mutual information 

Feature selection is a critical step in machine learning that involves selecting a subset of relevant features from a larger set of features in a dataset. This process is essential for improving model performance, reducing overfitting, and improving model interpretability.

In cases where the number of features is greater than the number of data points, such as in our dataset, feature selection becomes even more important.

While LASSO and ridge regression are commonly used for feature selection, they rely on the assumption that the input and output data follow a normal distribution. However, as we saw in Section 2.7, our data is far from normal. Therefore, in this chapter, we will use **mutual information**, a non-parametric measure of the dependence between two variables that does not assume any specific distribution, **for feature selection**.


As we need to predict a total of **four UPDRS scores**, we will calculate the mutual information **between each UPDRS score (y) and all features (X)**, and then use an **average ranking** approach to identify features that are highly dependent with Y.

# %% [code]
df_targets = df_with_CSF[["updrs_1","updrs_2","updrs_3","updrs_4"]]

from sklearn.feature_selection import mutual_info_regression

df_features = pd.read_csv("df_features_scaled_imputed.csv")	
df_features = df_features.drop(df_features.filter(regex='Unnamed').columns, axis=1)

dic = {}
for target in ["updrs_1","updrs_2","updrs_3","updrs_4"]:
	dic[target] = mutual_info_regression(df_features, df_targets[target], random_state = 42)

df_mi = pd.DataFrame(dic)
df_mi["feature_name"] = df_features.columns
ranks_df = df_mi[['updrs_1', 'updrs_2', 'updrs_3', 'updrs_4']].rank(axis=0, method='average',ascending=False)
df_mi['rank_sum'] = ranks_df.sum(axis=1)
feature_selected = df_mi[df_mi['rank_sum']  < 1450]['feature_name'].tolist() 
df_features[feature_selected].to_csv("df_features_scaled_imputed_selected.csv", index=False)

# %% [code]
#df_features_imputed_scaled_selected.csv
#!gdown --id 1dumKniqtezesFNoTfWRBln6gvdlPpbik
#the following features are selected
#selected_pro = ["O00533","O94919","P01009","P01857","P01860","P02787","P04433","P05060","P05090","P05452","P06454","P06727","P13591","P13611","P13987","P14618","P16035","P16152","P19021","P39060","P40925","P51884","P61916","Q06481","Q08380","Q12907","Q92520","Q92823","Q96KN2","Q99674","Q9BY67","Q9NQ79","Q9UNU6","Q99829"]
#selected_pep = ["GNPEPTFSWTK","IEIPSSVQQVPTIIK","VIAVNEVGR","VNGSPVDNHPFAGDVVFPR","AYQGVAAPFPK","ALMSPAGMLR","ILEVVNQIQDEER","MGNFPWQVFTNIHGR","NWGLGGHAFC(UniMod_4)R","DIPMNPMC(UniMod_4)IYR","NSPLDEENLTQENQDR","ANAGKPKDPTFIPAPIQAK","TVAAPSVFIFPPSDEQLK","TPLGDTTHTC(UniMod_4)PR","DASGVTFTWTPSSGK","DEPPQSPWDR","VEPLRAELQEGAR","AKLEEQAQQIR","LEEQAQQIR","LGADMEDVC(UniMod_4)GR","LQAEAFQAR","VQAAVGTSAAPVPSDNH","WELALGR","ATFGC(UniMod_4)HDGYSLDGPEEIEC(UniMod_4)TK","VC(UniMod_4)PFAGILENGAVR","ITYGETGGNSPVQEFTVPGSK","RPGGEPSPEGTTGQSYNQYSQR","SYTITGLQPGTDYK","FSGTWYAMAK","SDVVYTDWKK","GSPAINVAVHVFRK","C(UniMod_4)C(UniMod_4)TESLVNR","VHTEC(UniMod_4)C(UniMod_4)HGDLLEC(UniMod_4)ADDRADLAK","LC(UniMod_4)MAALK","DLLFRDDTVC(UniMod_4)LAK","HQTVPQNTGGKNPDPWAK","NFPSPVDAAFR","IYISGMAPR","LQSIGTENTEENRR","SSGLVSNAPGVQIR","GEAGAPGEEDIQGPTK","GYPGVQAPEDLEWER","SSQGGSLPSEEK","ISYGNDALMPSLTETK","LLDSLPSDTR","NSVIKVPMMNSK","C(UniMod_4)RDQLPYIC(UniMod_4)QFGIV","AGALNSNDAFVLK","EVVEEAENGR","LSPEDYTLK","VSTLPAITLK","C(UniMod_4)KPVNTFVHEPLVDVQNVC(UniMod_4)FQEK","NANTFISPQQR","EDC(UniMod_4)NELPPRR","RLEAGDHPVELLAR","GLSAEPGWQAK","RLEGQEEEEDNRDSSMK","YPGPQAEGDSEGLSQGLVDREK","KTLLSNLEEAK","QQTHMLDVMQDHFSR","TLLSNLEEAK","TLLSNLEEAKK","VAVVQYSGTGQQRPER","ALSSEWKPEIR","LRENELTYYC(UniMod_4)C(UniMod_4)KK","SC(UniMod_4)SPELQQK","GSGGLNLGNFFASR","MVQEQC(UniMod_4)C(UniMod_4)HSQLEELHC(UniMod_4)ATGISLANEQDR","RGYQLSDVDGVTC(UniMod_4)EDIDEC(UniMod_4)ALPTGGHIC(UniMod_4)SYR","LQDLYSIVR","FVEGLPINDFSR","AAFGQGSGPIMLDEVQC(UniMod_4)TGTEASLADC(UniMod_4)K","NLHGDGIALWYTR","GLVSWGNIPC(UniMod_4)GSK","HVEPGEPLAPSPQEPQAVGR","VHKEDDGVPVIC(UniMod_4)QVEHPAVTGNLQTQR","KNM(UniMod_35)FEFLK","NPDSSTTGPWC(UniMod_4)YTTDPTVR","QALPQVR"]


# %% [markdown]
### 3.2. Onehot encoding `upd23b_clinical_state_on_medication` and mergning all the data 

**`upd23b_clinical_state_on_medication` had three categories: On, Off, and NaN (No).** NaN indicates that Levodopa was not administered, On indicates a positive effect after administration, and Off indicates an ineffective result even after administration. To facilitate analysis, this was **transformed into three separate columns using One-hot encoding**. Additionally, the column name for `upd23b_clinical_state_on_medication` was modified to `upd23b` for convenience.


# %% [code]
df_with_CSF = df_with_CSF.drop(df_with_CSF.filter(regex='Unnamed').columns, axis=1)

  
#One hot encoding 'upd23b_clinical_state_on_medication'
df_with_CSF.rename(columns={'upd23b_clinical_state_on_medication':'upd23b'}, inplace=True)
df_with_CSF['upd23b'] = df_with_CSF['upd23b'].fillna(value="No")
df_with_CSF = pd.get_dummies(df_with_CSF, columns = ['upd23b'])


column_to_move = list(df_with_CSF.columns[:7]) + list(df_with_CSF.columns[-3:])
df_with_CSF = df_with_CSF[column_to_move]


df_features = pd.read_csv("df_features_scaled_imputed_selected.csv")
df_features = df_features.drop(df_features.filter(regex='Unnamed').columns, axis=1)

df_with_CSF['index_num'] = range(len(df_with_CSF))
df_features['index_num'] = range(len(df_features))

new_df = pd.concat([df_with_CSF.set_index('index_num'), df_features.set_index('index_num')], axis=1)

new_df.to_csv('integrated_dataset_scaled_imputed_selected.csv',index = False)


df_features = pd.read_csv("df_features_scaled_imputed.csv")
df_features = df_features.drop(df_features.filter(regex='Unnamed').columns, axis=1)

df_with_CSF['index_num'] = range(len(df_with_CSF))
df_features['index_num'] = range(len(df_features))

new_df = pd.concat([df_with_CSF.set_index('index_num'), df_features.set_index('index_num')], axis=1)

new_df.to_csv('integrated_dataset_scaled_imputed.csv',index = False)


# %% [code]
#integrated_dataset_imputed_scaled_selected.csv
#!gdown --id 1qaPcdcY51MCM629DaORp-4iwN_M8W1hp

# %% [markdown]
### 3.3. Dataset dividing for time series

Our ultimate objective is to develop a model that accurately predicts the updrs score at 0 months, 6 months, 12 months, and 24 months. To achieve this, we must create a new dataset that combines the current X with Y at 6 months, 12 months, and 24 months, respectively. The following code snippet makes this process possible. The code execution time takes roughly 5 to 10 minutes. If necessary, you may use the download link provided below to expedite the process.


# %% [code]

import warnings
warnings.filterwarnings('ignore')

df_final = pd.read_csv('integrated_dataset_scaled_imputed_selected.csv')
df_final.to_csv('t_plus_0.csv', index=False)

df_protein = pd.read_csv("/kaggle/input/amp-parkinsons-disease-progression-prediction/train_proteins.csv")
df_pebtide = pd.read_csv("/kaggle/input/amp-parkinsons-disease-progression-prediction/train_peptides.csv")

uniprot_list = list(df_protein["UniProt"].unique())
peptide_list = list(df_pebtide["Peptide"].unique())
patient_id_list = list(df_final["patient_id"].unique())


total_list= list(df_final.columns)
target_times = {6:pd.DataFrame(columns=total_list),
				12:pd.DataFrame(columns=total_list),
				24:pd.DataFrame(columns=total_list)}


for idx, patient_id in enumerate(patient_id_list):
	if (idx % 50 ==0):
		print(idx, len(patient_id_list))
	df_patient = df_final[df_final['patient_id'] ==patient_id]

	#print(list(df_patient["visit_id"].unique()))
	times = list(df_patient["visit_id"].unique())

	for si in range(len(times)):
		sid = times[si]
		st = int(times[si].split("_")[-1])

		for ei in range(si+1, len(times)):
			eid = times[ei]
			et = int(times[ei].split("_")[-1])

			if et-st not in target_times:
				continue	
			new_row = {}
			
			idx = total_list.index('updrs_4')
			X = df_final[df_final['visit_id'] ==sid][total_list[idx+1:]]
			Y = df_final[df_final['visit_id'] ==eid][total_list[:idx+1]]
			new_row.update(X.iloc[0].to_dict())
			new_row.update(Y.iloc[0].to_dict())
			target_times[et - st] = target_times[et - st].append(new_row, ignore_index=True)
			#target_times[et - st] = pd.concat(target_times[et - st], pd.Series(new_row))


target_times[6].to_csv("t_plus_6.csv", index = False)
target_times[12].to_csv("t_plus_12.csv", index = False)
target_times[24].to_csv("t_plus_24.csv", index = False)


# %% [markdown]
## 4. Training and evaluation 

In this section, we will train the model and see the results. If you want to skip steps 1-3 above and start training the model, you can start right here.


# %% [code]
#Downloading "elected feature versions"
#!gdown --id 1hsiCAqQVWFLchOJIPa1XE11Om0UVfBpA
#!gdown --id 1FtaSHoMD_ue-6f3p9iqARPEPHENsvpdd
#!gdown --id 1_ZXmmZiyC7QjXPMuTb-ukmqCeG05gnI_
#!gdown --id 1jz8rYm9Q6cXPXIU5_nOrdTBRzYXFwzcD

#!mkdir "dataset_cleaned"
#!mv t_plus_0_selected.csv ./dataset_cleaned/t_plus_0.csv
#!mv t_plus_6_selected.csv ./dataset_cleaned/t_plus_6.csv
#!mv t_plus_12_selected.csv ./dataset_cleaned/t_plus_12.csv
#!mv t_plus_24_selected.csv ./dataset_cleaned/t_plus_24.csv


#Downloading "all feature versions"
#!gdown --id 1G5PfjNgDvSS3FC-Zz1bkJc8A9GvJAp7F
#!gdown --id 1X9rJThgxsG7S2RgYf2nmPia0zFHyu6k8
#!gdown --id 11b6jkE4kkFiwAiwyV92Zen3PhTbfUw8H
#!gdown --id 1-kkUTaXH7Pa4pn5UOqHXhPCuLASaHWTJ

!mkdir "dataset_cleaned"
!mv t_plus_0.csv ./dataset_cleaned/t_plus_0.csv
!mv t_plus_6.csv ./dataset_cleaned/t_plus_6.csv
!mv t_plus_12.csv ./dataset_cleaned/t_plus_12.csv
!mv t_plus_24.csv ./dataset_cleaned/t_plus_24.csv

# %% [markdown]
### 4.1. Dataset checking before training


The current dataset we have created so far is t_plus_0, which has undergone feature selection. However, **t_plus_6 has only 137 data points**. Upon further examination, it was discovered that the **CSF mass spectrometry test is typically conducted annually**, resulting in a scarcity of data for a 6-month cycle.


| Time Difference | Rows | Columns |
| --- | --- | --- |
| t_plus_0 | 1068 | 139 |
| t_plus_6 | 137 | 139 |
| t_plus_12 | 543 | 139 |
| t_plus_24 | 435 | 139 |


# %% [code]
t_plus_0 = pd.read_csv("./dataset_cleaned/t_plus_0.csv")
t_plus_6 = pd.read_csv("./dataset_cleaned/t_plus_6.csv")
t_plus_12 = pd.read_csv("./dataset_cleaned/t_plus_12.csv")
t_plus_24 = pd.read_csv("./dataset_cleaned/t_plus_24.csv")

display(t_plus_0)
display(t_plus_6)
display(t_plus_12)
display(t_plus_24)

# %% [markdown]
### 4.2. Training models with hyperparamter cross validation

# %% [markdown]


After creating the data, it's time to train the regression models. This chapter covers the training and testing of **five regression models to predict the target** variable:

-   LinearRegression
-   DecisionTreeRegressor
-   ElasticNet
-   Bayesian Ridge
-   GradientBoostingRegressor

In addition to model training, this chapter also covers `hyperparameter search`, a technique used to determine the optimal performance of a model. To perform hyperparameter search, we'll use `GridSearchCV` from the `Scikit-learn` Python package. `GridSearchCV` uses `cross-validation` (CV) and `grid search` techniques to find the optimal hyperparameters. For a detailed explanation of these terms, please refer to the section below.

While training a model typically takes between 1 to 3 seconds, GridSearch may take longer as it tests a large number of hyperparameter combinations through cross-validation. In my case, using all **12 core 24 thread CPUs**, it took around **40 minutes** to obtain the final result.

# %% [markdown]

#### Hyperparameters

Hyperparameters are pre-set configurations or settings of a machine learning model that cannot be learned from the data, unlike the model parameters. These configurations control the model's behavior and performance during training, and their selection impacts the accuracy and generalization ability of the model. Some examples of hyperparameters include:

1.  Learning rate: the rate at which the model updates its parameters during training.
2.  Number of hidden layers: the number of layers in a neural network used to extract features from input data.
3.  Regularization parameters: penalties applied to the model to prevent overfitting.
4.  Number of estimators: the number of trees in a random forest algorithm.

#### Cross-validation

`Cross-validation` is a statistical technique used to evaluate the performance of a predictive model by dividing a dataset into multiple subsets, or "folds," and testing the model's accuracy on each fold. The goal of cross-validation is to provide an estimate of how well the model will perform on new, unseen data. The most common type of cross-validation is k-fold cross-validation, where the dataset is split into k subsets of roughly equal size. The model is then trained on k-1 of the subsets and tested on the remaining subset. This process is repeated k times, with each subset used as the test set once, and the average performance across all folds is calculated.

#### GridSearch

`GridSearch` is a technique used to identify the optimal hyperparameters for a machine learning model. In GridSearch, a set of possible values for each hyperparameter is defined, and the algorithm creates a grid of all possible combinations of hyperparameters. The algorithm then evaluates each combination of hyperparameters using a performance metric, such as Symmetric Mean Absolute Percentage Error (SMAPE), and selects the combination that performs the best. For example, in a Decision Tree Regressor model, we could define a set of possible values for each hyperparameter, such as:

-   Maximum depth: 2, 4, 6, 8, 10
-   Minimum samples required to split a node: 2, 4, 6, 8, 10
-   Minimum samples required to be at a leaf node: 1, 2, 3, 4, 5

`GridSearchCV` might train a Decision Tree model with a maximum depth of 2, minimum samples required to split a node of 4, and minimum samples required to be at a leaf node of 2 and evaluate its SMAPE on the validation (test) set using cross-validation techniques. It would repeat this process for all possible combinations of hyperparameters and select the combination that performs the best.

# %% [code]

from sklearn.ensemble import GradientBoostingRegressor
from sklearn.linear_model import BayesianRidge,LinearRegression,ElasticNet, SGDRegressor
from sklearn.tree import DecisionTreeRegressor

from sklearn.metrics import make_scorer
from sklearn.model_selection import GridSearchCV

from sklearn.model_selection import KFold
import time


# %% [code]

## This will take around few hours
def get_cv_dictionary(df):
	patient_ids = df['patient_id'].unique()
	kf = KFold(n_splits=5, shuffle= True, random_state=42)
	dic = {}
	for idx, (train_index, test_index) in enumerate(kf.split(patient_ids)):
		patient_train = patient_ids[train_index]
		patient_test = patient_ids[test_index]

		df_train_idx = np.where(df['patient_id'].isin(patient_train))[0]
		df_test_idx = np.where(df['patient_id'].isin(patient_test))[0]
		dic[idx] = (df_train_idx, df_test_idx)
	return dic

def custum_cv(X):
	for i in custom_cv_dic:
		yield custom_cv_dic[i]	

def run_training(f_out, df_y, df_X, updrs, file_name):

	models = [LinearRegression(),DecisionTreeRegressor(random_state=42),ElasticNet(random_state=42),BayesianRidge(),GradientBoostingRegressor(random_state=42)]

	def smape(y_true, y_pred):
		return np.mean((np.abs(y_true - y_pred) / (np.abs(y_true) + np.abs(y_pred)))) * 200

	# define the evaluation function to use SMAPE as scoring
	smape_scorer = make_scorer(smape, greater_is_better=False)

	param_grid = {
		"GradientBoostingRegressor": {'learning_rate': [0.01, 0.1, 1],'n_estimators': [50, 100, 150],'max_depth': [3, 5, 7],'min_samples_split': [2, 5, 10],'min_samples_leaf': [1, 2, 4]},
	 	"ElasticNet": {'alpha': [0.01, 0.1, 1.0],'l1_ratio': [0.1, 0.5, 0.9],'max_iter': [10000, 15000]},
	 	"SGDRegressor":{'loss': ['squared_loss', 'huber', 'epsilon_insensitive'],'penalty': ['none', 'l1', 'l2'],'alpha': [ 0.001, 0.01, 0.1],'l1_ratio': [0.3, 0.5, 0.7],'max_iter': [1000, 5000],'tol': [1e-3, 1e-4]},
	 	"BayesianRidge": {'alpha_1': [1e-6, 1e-5, 1e-4],'alpha_2': [1e-6, 1e-5, 1e-4],'lambda_1': [1e-6, 1e-5, 1e-4],'lambda_2': [1e-6, 1e-5, 1e-4],'tol': [1e-3, 1e-4, 1e-5],'n_iter': [100, 300, 500]},
		"DecisionTreeRegressor":{'max_depth': [2, 5, 10, 20, None],'min_samples_split': [2, 5, 10],'min_samples_leaf': [1, 2, 4],'max_features': ['auto', 'sqrt', 'log2']},
	 	"LinearRegression":{}}

	for model in models:
		st_time = time.time()
		modal_name = str(model).split("(")[0]
		# create the grid search object
		grid_search = GridSearchCV(estimator=model, param_grid=param_grid[modal_name], cv=list(custom_cv_dic.values()), scoring=smape_scorer, n_jobs=-1)

		# fit the grid search to the training data
		grid_search.fit(df_X, df_y)

		# The score may have a negative value due to the GridSearch pipeline problems, but can be replaced with positive numbers.
		took_time = time.time()- st_time
		str_out = file_name+","+updrs+","+modal_name+","+str(-grid_search.best_score_)+ "," + str(grid_search.best_params_).replace(",","|")+","+str(took_time)
		f_out.write(str_out+"\n")
		print(str_out)

updrs_list = ['updrs_1','updrs_2','updrs_3','updrs_4']
f_out = open("training_result.csv","a")
f_out.write("file_name,updrs,model_name,SMAPE,hyperparameter,run_time\n")

file_names = os.listdir("dataset_cleaned")

for dataset_path in file_names:
	print(dataset_path)
	df = pd.read_csv("./dataset_cleaned/"+dataset_path)
	global custom_cv_dic 
	custom_cv_dic = get_cv_dictionary(df)

	for updrs in updrs_list:
		idx = list(df.columns).index("updrs_4")+1
		df_y = df[updrs]
		df_X = df[df.columns[idx:]]
		run_training(f_out,df_y, df_X,updrs, dataset_path)


# %% [markdown]
### 4.3. Reading results

In the test set, we need to make 16 predictions across two axes. The first axis comprises four different time intervals, while the second axis comprises four different UPDRS scores. For each time interval and UPDRS score combination, we have prepared five hyperparameter-tuned models. Consequently, the "training_result.csv" file contains 80 rows, each representing a combination of **four time intervals**, **four UPDRS scores**, and **five model variants**.

Each row in the file provides the 1) time interval, 2) UPDRS score type, 3) SMAPE score of the model, 4) hyperparameter information of the model, and the 5) total training time. Since we only require the model with the lowest SMAPE score out of the 16 predictions, the final outcome can be presented in the table below.


#### Key Findings:
1. Predicting UPDRS 1 is generally easiest.
2. UPDRS 4 is very difficult to predict, with high SMAPE scores across all time intervals.
3. Despite the small dataset, t+6 had the lowest SMAPE score.
4. Gradient boosting works well for predicting UPDRS 1-3.
5. Decision trees are most effective for UPDRS 4 prediction.
6. GBR takes the longest to train, but performs well among the five models.

#### Discussion:
1. **High SMAPE scores for UPDRS 4**: may be due to the metric's **greater penalty on high predictions**. Since **imputation** filled 499/1068 missing values **with 0**, around **half of the y_true** will have the **lowest value**. Then most predictions over 0 will have a maximum value of 200 SMAPE. If imputation was performed with a non-zero mean or median value, SMAPE would be significantly reduced. Alternatively, if the contest staff filled missing values of UPDRS 4 with 0 in the test set, our imputation assumption is not wrong, so the high SMAPE score is likely due to dataset characteristics.

2. **T+6 prediction**: **CSF mass spectrometry** was primarily conducted on an **annual basis**, as mentioned in Section 4.1. Therefore, the "t_plus_6.csv" dataset contains fewer samples than the others, with only 123 rows. Despite this, the SMAPE score was impressively low. Since cross-validation was transparent, the training process had no issues, indicating no overfitting. Therefore, it appears that the **"t_plus_6"** dataset **may have commonalities that make it easier to predict**. Further investigation is needed to confirm this possibility.


# %% [code]
#Downloding "training_result.csv"
#!gdown --id "1TBW4eX-TqEas16MyqFrSDDJPOftHVchd"
df_result = pd.read_csv("training_result.csv")
display(df_result)

# %% [markdown]
| T plus | Features | updrs_1 | updrs_2 | updrs_3 | updrs_4 |
|--------|----------------------------|---------|---------|---------|---------|
| 0      | Selected (131 + 3 Levodopa)| 65.86   | 94.14   | 86.65   | 158.10  |
| 6      | Selected (131 + 3 Levodopa)| 60.18   | 59.20   | 53.07   | 185.07  |
| 12     | Selected (131 + 3 Levodopa)| 64.69   | 93.92   | 88.42   | 157.07  |
| 24     | Selected (131 + 3 Levodopa)| 65.45   | 94.78   | 90.95   | 156.90  |


| T plus | Features| updrs_1                   | updrs_2                  | updrs_3                   | updrs_4               |
|--------|----------------------------|---------------------------|--------------------------|---------------------------|-----------------------|
| 0      | selected (131 + 3 Levodopa)| DecisionTreeRegressor     |GradientBoostingRegressor | ElasticNet | DecisionTreeRegressor |
| 6      | selected (131 + 3 Levodopa)| GradientBoostingRegressor |ElasticNet                | ElasticNet | DecisionTreeRegressor |
| 12     | selected (131 + 3 Levodopa)| GradientBoostingRegressor |GradientBoostingRegressor | ElasticNet | DecisionTreeRegressor |
| 24     | selected (131 + 3 Levodopa) | GradientBoostingRegressor |ElasticNet                | ElasticNet | DecisionTreeRegressor |

			

# %% [code]
df_result = pd.read_csv("training_result.csv")
mix_SMAPE_row = df_result.loc[df_result.groupby(['file_name', 'updrs'])['SMAPE'].idxmin()]
mix_SMAPE_row.to_csv("result_cleaned.csv", index=False)

df_result.columns

plt.subplots(figsize=(16,5),constrained_layout=True)

plt.subplot(1, 2, 1)    
ax = sns.countplot(x=mix_SMAPE_row["model_name"])
ax.set_title("# of best selected models")

plt.subplot(1,2, 2)    
ax = sns.boxplot(data = df_result, x="model_name", y="run_time")
ax.set_title("Runding time per one training")
ax.set_xticklabels(["LR", "DTR", "EN", "BR", "GBR" ])


# %% [markdown]
### 4.4. Setting final models

Now, let's create a model with the best hyperparameter based on 'training_result.csv' and train it with the full dataset.


# %% [code]

df = pd.read_csv('training_result.csv')
max_SMAPE_row = df.loc[df.groupby(['file_name', 'updrs'])['SMAPE'].idxmin()]
max_SMAPE_row.to_csv("result_cleaned.csv", index=False)

file_dic = {"t_plus_0.csv":"t_plus_0","t_plus_6.csv":"t_plus_6","t_plus_12.csv":"t_plus_12","t_plus_24.csv":"t_plus_24"}
models = {"LinearRegression":LinearRegression(),"DecisionTreeRegressor":DecisionTreeRegressor(random_state=42),"ElasticNet":ElasticNet(random_state=42),"BayesianRidge":BayesianRidge(),"GradientBoostingRegressor":GradientBoostingRegressor(random_state=42)}


model_dic = {
"t_plus_0": {"updrs_1":None,"updrs_2":None,"updrs_3":None,"updrs_4":None},
"t_plus_6": {"updrs_1":None,"updrs_2":None,"updrs_3":None,"updrs_4":None},
"t_plus_12":{"updrs_1":None,"updrs_2":None,"updrs_3":None,"updrs_4":None},
"t_plus_24":{"updrs_1":None,"updrs_2":None,"updrs_3":None,"updrs_4":None}}


for row in max_SMAPE_row.iterrows():
    file_name = row[1]["file_name"]
    updrs = row[1]["updrs"]
    hyperparameter = eval(row[1]["hyperparameter"].replace("|",","))
    run_time = row[1]["run_time"]
    model_name = row[1]["model_name"]

    model = models[model_name]
    model.__init__(**hyperparameter)

    df_total = pd.read_csv("./dataset_cleaned/"+ file_name)

    idx = list(df_total.columns).index("updrs_4")+1
    df_y = df_total[updrs]
    df_X = df_total[df_total.columns[idx:]]
    model.fit(df_X,df_y)

    model_dic[file_dic[file_name]][updrs] = {"model":models[model_name].__init__(**hyperparameter), "X": df_X, "y":df_y}
pickle.dump(model_dic, open("prediction_models.pkl", "wb"))


# %% [code]
##Downloading "prediction_models.pkl"
#!gdown --id "1te1gz7zoPCbtrJMkf_XZDLJ44BAZgfY-"


# %% [markdown]
## 5. Finding candidate biomarkers

In this section, we will identify the proteins or peptides that have the greatest impact on the final predictions. Typically, we utilize the SHapley Additive exPlanations (SHAP) method for interpretability, which assumes independence and additivity between features. However, in our specific case, this assumption may not hold perfectly due to the possibility of a peptide being present in multiple proteins. To address this challenge, we employ two additional interpretability methods: permutation feature importance (PFI) and built-in feature importance.

PFI evaluates the significance of each feature by permuting their values and measuring the resulting effect on the model's performance. By comparing the model's performance before and after permutation, we can calculate a PFI score for each feature. Higher PFI scores indicate greater importance, as permuting an influential feature leads to a more notable decrease in performance. This method takes into account feature interactions and non-linear relationships, providing a reliable measure of feature importance.

Conversely, the built-in feature importance relies on the inherent interpretability of the model itself. For instance, in a regression model, the coefficients can indicate the importance of each feature. Accessing this built-in feature importance can be achieved using the feature_importance method in models implemented with scikit-learn.

By employing the SHAP, PFI, and built-in feature importance techniques, we can gain valuable insights into the contributions of individual protein and peptides within our models. This information enables us to make discover the candidate biomarkers for Parkinson's diseace as well.


# %% [markdown]
### 5.1. Calculating three different interpretable AI methods 


# %% [code]

#model_dic = pickle.load(open('prediction_models.pkl',"rb"))
column_names = ["upd23b_No","upd23b_Off","upd23b_On","O00533","O94919","P01009","P01019","P01857","P01860","P02787","P04156","P04433","P05060","P05452","P05546","P06454","P06727","P08603","P13591","P13987","P14618","P16035","P16152","P39060","P51884","P61916","P80748","Q06481","Q08380","Q12907","Q13332","Q92520","Q92823","Q96KN2","Q96PD5","Q99674","Q9BY67","Q9NQ79","Q99829","GNPEPTFSWTK","IEIPSSVQQVPTIIK","SMEQNGPGLEYR","VIAVNEVGR","VNGSPVDNHPFAGDVVFPR","AYQGVAAPFPK","ILEVVNQIQDEER","MGNFPWQVFTNIHGR","HSIFTPETNPR","NWGLGGHAFC(UniMod_4)R","DIPMNPMC(UniMod_4)IYR","LVDKFLEDVKK","ISLPESLK","TVAAPSVFIFPPSDEQLK","TPLGDTTHTC(UniMod_4)PR","DASGVTFTWTPSSGK","NFPPSQDASGDLYTTSSQLTLPATQC(UniMod_4)LAGK","VEPLRAELQEGAR","AKLEEQAQQIR","LEEQAQQIR","LGADMEDVC(UniMod_4)GR","LQAEAFQAR","VQAAVGTSAAPVPSDNH","WELALGR","GGSTSYGTGSETESPR","C(UniMod_4)SYTEDAQC(UniMod_4)IDGTIEVPK","VC(UniMod_4)PFAGILENGAVR","ITYGETGGNSPVQEFTVPGSK","NLQPASEYTVSLVAIKGNQESPK","RPGGEPSPEGTTGQSYNQYSQR","SYTITGLQPGTDYK","FSGTWYAMAK","SDVVYTDWKK","GSPAINVAVHVFRK","C(UniMod_4)C(UniMod_4)TESLVNR","KQTALVELVK","VHTEC(UniMod_4)C(UniMod_4)HGDLLEC(UniMod_4)ADDRADLAK","LC(UniMod_4)MAALK","DLLFRDDTVC(UniMod_4)LAK","IMNGEADAMSLDGGFVYIAGK","NFPSPVDAAFR","IYISGMAPR","LQSIGTENTEENRR","VVEQMC(UniMod_4)ITQYER","LEPGQQEEYYR","SSGLVSNAPGVQIR","GYPGVQAPEDLEWER","SSQGGSLPSEEK","ISYGNDALMPSLTETK","STNLHDYGMLLPC(UniMod_4)GIDK","WYFDVTEGK","C(UniMod_4)PNPPVQENFDVNK","LLDSLPSDTR","NSVIKVPMMNSK","C(UniMod_4)RDQLPYIC(UniMod_4)QFGIV","AGALNSNDAFVLK","AGKEPGLQIWR","EVVEEAENGR","VSTLPAITLK","C(UniMod_4)KPVNTFVHEPLVDVQNVC(UniMod_4)FQEK","VTGVVLFR","NANTFISPQQR","EDC(UniMod_4)NELPPRR","RLEAGDHPVELLAR","C(UniMod_4)FSGQC(UniMod_4)ISK","GLSAEPGWQAK","RLEGQEEEEDNRDSSMK","YPGPQAEGDSEGLSQGLVDREK","KTLLSNLEEAK","TLLSNLEEAK","TLLSNLEEAKK","LRENELTYYC(UniMod_4)C(UniMod_4)KK","AVSEKEVDSGNDIYGNPIKR","SC(UniMod_4)SPELQQK","SNAQGIDLNR","SSALDMENFR","MVQEQC(UniMod_4)C(UniMod_4)HSQLEELHC(UniMod_4)ATGISLANEQDR","RGYQLSDVDGVTC(UniMod_4)EDIDEC(UniMod_4)ALPTGGHIC(UniMod_4)SYR","QKWEAEPVYVQR","LQDLYSIVR","FSGSNSGNTATLTISR","AAFGQGSGPIMLDEVQC(UniMod_4)TGTEASLADC(UniMod_4)K","NLHGDGIALWYTR","SSNTYTLTDVRR","SRYPVC(UniMod_4)GSDGTTYPSGC(UniMod_4)QLR","LVWEEAMSR","LIADLGSTSITNLGFR","VFNTPEGVPSAPSSLK","GLVSWGNIPC(UniMod_4)GSK","HVEPGEPLAPSPQEPQAVGR","VHKEDDGVPVIC(UniMod_4)QVEHPAVTGNLQTQR","NPDSSTTGPWC(UniMod_4)YTTDPTVR","QALPQVR"]
                
f_out = open("feature_importances.csv","w")
f_out.write("file_name,updrs,importance_type,feature_name,importance_mean,importance_std\n")

# Calculate built-in feature importances
for file_name in model_dic:
    for updrs in model_dic[file_name]:
        model = model_dic[file_name][updrs]['model']
        df_X = model_dic[file_name][updrs]['X']
        df_y = model_dic[file_name][updrs]['y']

        model.fit(df_X,df_y)
        

        if hasattr(model, 'feature_importances_'):
            for name, val in zip(column_names, model.feature_importances_):
                f_out.write(file_name+","+updrs+","+"tree"+","+ name+","+str(val)+","+str(0)+"\n")
        else:
            f_out.write(file_name+","+updrs+","+"tree"+","+ name+","+str(0)+","+str(0)+"\n")

f_out.close()

# Calculate permutation feature importance
from sklearn.inspection import permutation_importance

f_out = open("feature_importances.csv","a")

for file_name in model_dic:
    for updrs in model_dic[file_name]:
        model = model_dic[file_name][updrs]['model']
        df_X = model_dic[file_name][updrs]['X']
        df_y = model_dic[file_name][updrs]['y']

        model.fit(df_X,df_y)

        r = permutation_importance(model, df_X, df_y,n_repeats=30,random_state=0)

        for name, mean, std in zip(column_names, r.importances_mean,r.importances_std):
            f_out.write(file_name+","+updrs+","+"permutation feature impoartance"+","+ name+","+str(mean)+","+str(std)+"\n")

f_out.close()


# Calculate SHAP feature importance
import shap

f_out = open("feature_importances.csv","a")

import numpy as np
for file_name in model_dic:
    for updrs in model_dic[file_name]:
        model = model_dic[file_name][updrs]['model']
        df_X = model_dic[file_name][updrs]['X']
        df_y = model_dic[file_name][updrs]['y']

        model.fit(df_X,df_y)

        importances_mean = np.zeros((132))
        try:           
            explainer = shap.Explainer(model)
            shap_values = explainer(df_X)
            importances_mean = abs(shap_values.values).sum(axis = 0)
            model_dic[file_name][updrs]['shap_values'] = shap_values
        except:
            pass

        for name, mean in zip(column_names, importances_mean):
            f_out.write(file_name+","+updrs+","+"SHAP"+","+ name+","+str(mean)+","+str(0)+"\n")

f_out.close()
pickle.dump(model_dic, open("prediction_models_with_shap.pkl", "wb"))


# %% [code]
#Downloading feature importances
#!gdown --id "1o-oP-u_EgzBQxrefeqctGnfLsKok95LL"


# %% [code]
#Plotting results

df = pd.read_csv("feature_importances.csv")

for time_diff in ["t_plus_0","t_plus_6","t_plus_12","t_plus_24"]:
    for updrs in ['updrs_1','updrs_2','updrs_3','updrs_4']:
        plt.subplots(figsize=(16,5),constrained_layout=True)

        for idx, importance_type in enumerate(["Built-in","Permutation feature impoartance","SHAP"]):

            df_pick = df[(df['file_name'] ==time_diff) & (df['updrs'] ==updrs) & (df['importance_type'] ==importance_type)]
            df_sorted = df_pick.sort_values('importance_mean', ascending=False)

            plt.subplot(1, 3, idx+1)    
            ax = sns.barplot(data = df_sorted.iloc[:10], x = "feature_name", y= "importance_mean")
            print(time_diff + ", " + updrs + ", " + importance_type,"|".join(list(df_sorted.iloc[:4]["feature_name"])))
            ax.set_title(time_diff + ", " + updrs + ", " + importance_type )
            ax.tick_params(axis='x', rotation=90)


# %% [markdown]
### 5.2 Summarizing the all results


In the previous analysis, we observed variations in the results across different time intervals and UPDRS scores. Notably, Levodopa-related features consistently emerged as the most important, except for the T+6 interval. To avoid overwhelming complexity in the figures, we aim to summarize the results.

However, a challenge arises due to the disparate scales of importance values for each interpretability method. For instance, SHAP values can range in the thousands, while PFI values typically remain below 1. Therefore, a consolidation of the results becomes necessary.

To address this challenge, we propose utilizing a ranking-based synthesis method. We will sum the importance values from each interpretability method and generate three rankings in descending order based on the highest value. Subsequently, we will compute the average value of these rankings to obtain the final ranking. This approach allows us to effectively merge the information from the three interpretability methods, providing a comprehensive overview of feature importance in a more concise manner.


# %% [code]
df = pd.read_csv("feature_importances.csv")
['file_name', 'updrs', 'importance_type', 'feature_name', 'importance_mean', 'importance_std']
df = df[df['updrs'] != 'updrs_4']

df_b = df[df['importance_type']=='built_in']
df_p = df[df['importance_type']=='permutation feature impoartance']
df_s = df[df['importance_type']=='SHAP']

summed_df_b = df_b.groupby('feature_name')['importance_mean'].sum()
summed_df_p = df_p.groupby('feature_name')['importance_mean'].sum()
summed_df_s = df_s.groupby('feature_name')['importance_mean'].sum()


# rank the values in each DataFrame
ranked_df_b = summed_df_b.rank(method='dense', ascending=False)
ranked_df_p = summed_df_p.rank(method='dense', ascending=False)
ranked_df_s = summed_df_s.rank(method='dense', ascending=False)


# merge the ranked DataFrames
merged_df = pd.concat([ranked_df_b, ranked_df_p, ranked_df_s], axis=1)
merged_df.columns = ['Built_in','PFI','SHAP']
merged_df['average_rank'] = merged_df.mean(axis=1)

sorted_df =merged_df.sort_values('average_rank')
sorted_df = merged_df.sort_values('average_rank')

sorted_df_b =summed_df_b.sort_values(ascending=False)
sorted_df_p =summed_df_p.sort_values(ascending=False)
sorted_df_s =summed_df_s.sort_values(ascending=False)

sorted_df.to_csv('importance_ranking.csv')

# %% [code]
#Downloading importances ranking
#!gdown --id "1q2aOC6iaf0d2ihpaw0urOJqA0CPqFgW6"

# %% [markdown]
With the exception of `upd23b_No`, the rankings exhibit irregularities. For instance, although `LQDLYSIVR` received high scores in SHAP (2nd) and PFI (3rd), it did not perform well in the Built-in (54th) method. This discrepancy can be attributed to the inclusion of models with low performance or limited data points, such as UPDRS4 or T+6, in the results. Additionally, differences in the calculation methods of the interpretability techniques could contribute to these variations. This further emphasizes the importance of not relying solely on a single method when determining candidate biomarkers.


| Feature_name                  | Built_in | PFI | SHAP | Average rank |
|-------------------------------|:--------:|:---:|:----:|:------------:|
| upd23b_No                     |     1    |  1  |   1  |     1.00     |
| Q06481                        |    10    |  5  |   3  |     6.00     |
| AVSEKEVDSGNDIYGNPIKR          |    12    |  6  |   4  |     7.33     |
| ISYGNDALMPSLTETK              |    11    |  11 |  10  |     10.67    |
| P04433                        |     7    |  8  |  18  |     11.00    |
| TPLGDTTHTC(UniMod_4)PR        |    35    |  2  |   7  |     14.67    |
| NLQPASEYTVSLVAIKGNQESPK       |    23    |  15 |  16  |     18.00    |
| LQDLYSIVR                     |    54    |  3  |   2  |     19.67    |
| FSGTWYAMAK                    |     8    |  12 |  45  |     21.67    |
| C(UniMod_4)FSGQC(UniMod_4)ISK |     6    |  24 |  35  |     21.67    |
| LIADLGSTSITNLGFR              |    19    |  25 |  22  |     22.00    |
| TLLSNLEEAKK                   |    45    |  14 |   8  |     22.33    |


The table provided below contains compiled information from Uniprot and Protein Data Bank. It includes 11 proteins and peptides associated with Alzheimer's disease, a brain disorder, as well as proteins associated with cancer. 

For more detailed information on the top 11 protein peptides, you can refer to the following links:
- Top 11 Protein Peptides summary and All ranks - [Google Sheets](https://docs.google.com/spreadsheets/d/1TAIsb41U-mO0Ot5ZYPxYIglemiErXkhodiW25JWI90E/edit?usp=sharing)
- Top 11 Protein Peptides detailed descriptions - [Google Docs](https://docs.google.com/document/d/1IKTu2hL0SFZJHU2AYKur6sSjRn5mKeJm/edit?usp=sharing&ouid=101633548124306174896&rtpof=true&sd=true)

For information on other proteins and peptides that are not part of the ranking, you can utilize our separate search code available at the following link: [Protein and Peptide Search Code - Google Colab](https://colab.research.google.com/drive/1EcqZmy0QL4QFJoIVa65RsttOQz9lxcg6?usp=sharing#scrollTo=sSNJplQ-iCw5).

| Feature name            | Uniprot ID | Description                           | Abbreviation | Alzheimer's Disease | Cancer |
|-------------------------|:----------:|---------------------------------------|:------------:|:-------------------:|:------:|
| Q06481                  |   Q06481   | Amyloid beta precursor like protein 2 |     APLP2    |          O          |    O   |
| AVSEKEVDSGNDIYGNPIKR    |   P16035   | Metalloproteinase inhibitor 2         |    TIMP-2    |          O          |    O   |
| ISYGNDALMPSLTETK        |   P05067   | Amyloid-beta precursor protein        |      APP     |          O          |    O   |
| P04433                  |   P04433   | Immunoglobulin kappa variable 3-11    |   IGKV3-11   |          X          |    O   |
| TPLGDTTHTCPR            |   P01860   | Immunoglobulin heavy constant gamma 3 |     IGHG3    |          O          |    O   |
| NLQPASEYTVSLVAIKGNQESPK |   P02751   | Fibronectin                           |      FN1     |          O          |    O   |
| LQDLYSIVR               |   P39060   | Collagen alpha-1(XVIII) chain         |    COL18A1   |          X          |    O   |
| FSGTWYAMAK              |   P02753   | Retinol-binding protein 4             |     RBP4     |          O          |    O   |
| CFSGQCISK               |   P10643   | Complement component C7               |      C7      |          O          |    O   |
| LIADLGSTSITNLGFR        |   Q92520   | Protein FAM3C                         |     FAM3C    |          O          |    O   |
| TLLSNLEEAKK             |   P10909   | Clusterin                             |      CLU     |          O          |    O   |
```
