## Supplementary material (6) Additional Pipeline Codes for "Discovering Biomarker Proteins and Peptides for Parkinson’s Disease Prognosis Prediction with Machine Learning and Interpretability Methods": model.py

```
import torch
from torch import nn
from torch.utils.data import Dataset
import pandas as pd
import code
# Define SMAPE loss function
def smape_loss(y_true, y_pred):
    return 100 * torch.mean(2 * torch.abs(y_true - y_pred) / (torch.abs(y_true) + torch.abs(y_pred)))

# Create custom Dataset class
class RegressionDataset(Dataset):
    def __init__(self, df, input_cols, output_cols):
        self.data = df
        self.input_cols = input_cols
        self.output_cols = output_cols
    
    def __len__(self):
        return len(self.data)
    
    def __getitem__(self, idx):
        #code.interact(local=dict(globals(), **locals()))
        return self.data.iloc[idx][self.input_cols].values, self.data.iloc[idx][self.output_cols].values

# Create PyTorch Lightning model
class RegressionModel(torch.nn.Module):
    def __init__(self, input_dim, output_dim, layers, learning_rate=1e-3, dropout=0.5):
        super().__init__()
        self.layers = layers
        self.learning_rate = learning_rate
        self.dropout = nn.Dropout(dropout)

        self.model = nn.Sequential(
            nn.Linear(input_dim, layers[0]),
            nn.ReLU(),
            *[l for layer_pair in zip(layers[:-1], layers[1:]) for l in [nn.Linear(layer_pair[0], layer_pair[1]), nn.ReLU(), self.dropout]],
            nn.Linear(layers[-1], output_dim)
        )

    def forward(self, x):
        #code.interact(local=dict(globals(), **locals()))
        return self.model(x)        
```
