## Supplementary material (6) Additional Pipeline Codes for "Discovering Biomarker Proteins and Peptides for Parkinson’s Disease Prognosis Prediction with Machine Learning and Interpretability Methods": train.py

```
import pandas as pd
import torch
from torch.utils.data import DataLoader
import pytorch_lightning as pl
from pytorch_lightning import Trainer
from pytorch_lightning.callbacks import EarlyStopping, ModelCheckpoint, LearningRateMonitor
from pytorch_lightning.loggers import WandbLogger
from model import RegressionDataset, RegressionModel, smape_loss
from sklearn.model_selection import ParameterGrid, KFold
import torch.optim as optim

import numpy as np
import code, os
import random, time
class PLRegressionModel(pl.LightningModule):
    def __init__(self, input_dim, output_dim, layers, learning_rate=1e-3, dropout=0.5):
        super().__init__()
        self.save_hyperparameters()
        self.model = RegressionModel(input_dim,output_dim,layers,learning_rate,dropout)

    def forward(self, x):
        return self.model(x)

    def training_step(self, batch, batch_idx):
        x, y = batch
        y_pred = self.model(x)
        loss = smape_loss(y, y_pred)
        self.log("train_loss", loss)
        return loss

    def validation_step(self, batch, batch_idx):
        x, y = batch
        y_pred = self.model(x)
        loss = smape_loss(y, y_pred)
        self.log("val_loss", loss)

    def configure_optimizers(self):
        optimizer = torch.optim.Adam(self.parameters(), lr=self.hparams.learning_rate)
        scheduler = optim.lr_scheduler.ReduceLROnPlateau(optimizer, mode='min', factor=0.5, patience=5, verbose=True)
        return {"optimizer": optimizer, "lr_scheduler": scheduler, "monitor": "val_loss"}


def get_cv_dictionary(df):
    patient_ids = df['patient_id'].unique()
    kf = KFold(n_splits=5, shuffle= True, random_state=42)
    dic = {}
    for idx, (train_index, test_index) in enumerate(kf.split(patient_ids)):
        patient_train = patient_ids[train_index]
        patient_test = patient_ids[test_index]

        df_train_idx = np.where(df['patient_id'].isin(patient_train))[0]
        df_test_idx = np.where(df['patient_id'].isin(patient_test))[0]
        dic[idx] = (df_train_idx, df_test_idx)
    return dic


def get_dataloader(df, input_cols, output_cols, train_index, validation_index, batch_size=16):
    train_data = df.iloc[train_index]
    val_data = df.iloc[validation_index]

    train_dataset = RegressionDataset(train_data, input_cols, output_cols)
    val_dataset = RegressionDataset(val_data, input_cols, output_cols)
    
    train_loader = DataLoader(train_dataset, batch_size=batch_size, shuffle=True)
    val_loader = DataLoader(val_dataset, batch_size=batch_size, shuffle=False)
    
    return train_loader, val_loader

def train_model(train_data, input_cols, output_cols, model_params, cv, max_epochs=10, batch_size=16):
    param_grid = ParameterGrid(model_params)
    best_loss = np.inf


    for params in param_grid:
        val_losses = []

        for fold_idx, (train_idx, val_idx) in enumerate(cv):
            train_loader, val_loader = get_dataloader(train_data, input_cols, output_cols, train_idx, val_idx, batch_size=batch_size)

            # Callbacks
            early_stop = EarlyStopping(monitor="val_loss", patience=10)
            checkpoint = ModelCheckpoint(monitor="val_loss", mode="min",save_top_k=1)
            lr_logger = LearningRateMonitor(logging_interval="step")


            #train_data[train_idx]
            model = PLRegressionModel(len(input_cols), len(output_cols), **params)

            trainer = Trainer(
                max_epochs=max_epochs,
                gpus=1 if torch.cuda.is_available() else 0,
                callbacks=[early_stop, checkpoint, lr_logger],
            )

            trainer.fit(model, train_loader, val_loader)
            val_loss = trainer.logged_metrics['val_loss']
            val_losses.append(val_loss)        

        if np.mean(val_losses) < best_loss:
            best_loss = val_loss
            best_params = params
                
    return best_params,best_loss


# Set random seed for reproducibility
seed = 42
torch.manual_seed(seed)
torch.cuda.manual_seed_all(seed)
random.seed(seed)
np.random.seed(seed)


df_clinical = pd.read_csv("../train_clinical_data.csv")
df_protein = pd.read_csv("../train_proteins.csv")
df_pebtide = pd.read_csv("../train_peptides.csv")


df_total = pd.read_csv("../dataset_cleaned_mk2/t_plus_0.csv")
df_total = df_total.drop(df_total.filter(regex='Unnamed').columns, axis=1)


idx = list(df_total.columns).index("upd23b_On")+1
main_info_y_onehot = list(df_total.columns[:idx])
selected_pro = ["O00533","O94919","P01009","P01019","P01857","P01860","P02787","P04156","P04433","P05060","P05452","P05546","P06454","P06727","P08603","P13591","P13987","P14618","P16035","P16152","P39060","P51884","P61916","P80748","Q06481","Q08380","Q12907","Q13332","Q92520","Q92823","Q96KN2","Q96PD5","Q99674","Q9BY67","Q9NQ79","Q99829"]
selected_pep = ["GNPEPTFSWTK","IEIPSSVQQVPTIIK","SMEQNGPGLEYR","VIAVNEVGR","VNGSPVDNHPFAGDVVFPR","AYQGVAAPFPK","ILEVVNQIQDEER","MGNFPWQVFTNIHGR","HSIFTPETNPR","NWGLGGHAFC(UniMod_4)R","DIPMNPMC(UniMod_4)IYR","LVDKFLEDVKK","ISLPESLK","TVAAPSVFIFPPSDEQLK","TPLGDTTHTC(UniMod_4)PR","DASGVTFTWTPSSGK","NFPPSQDASGDLYTTSSQLTLPATQC(UniMod_4)LAGK","VEPLRAELQEGAR","AKLEEQAQQIR","LEEQAQQIR","LGADMEDVC(UniMod_4)GR","LQAEAFQAR","VQAAVGTSAAPVPSDNH","WELALGR","GGSTSYGTGSETESPR","C(UniMod_4)SYTEDAQC(UniMod_4)IDGTIEVPK","VC(UniMod_4)PFAGILENGAVR","ITYGETGGNSPVQEFTVPGSK","NLQPASEYTVSLVAIKGNQESPK","RPGGEPSPEGTTGQSYNQYSQR","SYTITGLQPGTDYK","FSGTWYAMAK","SDVVYTDWKK","GSPAINVAVHVFRK","C(UniMod_4)C(UniMod_4)TESLVNR","KQTALVELVK","VHTEC(UniMod_4)C(UniMod_4)HGDLLEC(UniMod_4)ADDRADLAK","LC(UniMod_4)MAALK","DLLFRDDTVC(UniMod_4)LAK","IMNGEADAMSLDGGFVYIAGK","NFPSPVDAAFR","IYISGMAPR","LQSIGTENTEENRR","VVEQMC(UniMod_4)ITQYER","LEPGQQEEYYR","SSGLVSNAPGVQIR","GYPGVQAPEDLEWER","SSQGGSLPSEEK","ISYGNDALMPSLTETK","STNLHDYGMLLPC(UniMod_4)GIDK","WYFDVTEGK","C(UniMod_4)PNPPVQENFDVNK","LLDSLPSDTR","NSVIKVPMMNSK","C(UniMod_4)RDQLPYIC(UniMod_4)QFGIV","AGALNSNDAFVLK","AGKEPGLQIWR","EVVEEAENGR","VSTLPAITLK","C(UniMod_4)KPVNTFVHEPLVDVQNVC(UniMod_4)FQEK","VTGVVLFR","NANTFISPQQR","EDC(UniMod_4)NELPPRR","RLEAGDHPVELLAR","C(UniMod_4)FSGQC(UniMod_4)ISK","GLSAEPGWQAK","RLEGQEEEEDNRDSSMK","YPGPQAEGDSEGLSQGLVDREK","KTLLSNLEEAK","TLLSNLEEAK","TLLSNLEEAKK","LRENELTYYC(UniMod_4)C(UniMod_4)KK","AVSEKEVDSGNDIYGNPIKR","SC(UniMod_4)SPELQQK","SNAQGIDLNR","SSALDMENFR","MVQEQC(UniMod_4)C(UniMod_4)HSQLEELHC(UniMod_4)ATGISLANEQDR","RGYQLSDVDGVTC(UniMod_4)EDIDEC(UniMod_4)ALPTGGHIC(UniMod_4)SYR","QKWEAEPVYVQR","LQDLYSIVR","FSGSNSGNTATLTISR","AAFGQGSGPIMLDEVQC(UniMod_4)TGTEASLADC(UniMod_4)K","NLHGDGIALWYTR","SSNTYTLTDVRR","SRYPVC(UniMod_4)GSDGTTYPSGC(UniMod_4)QLR","LVWEEAMSR","LIADLGSTSITNLGFR","VFNTPEGVPSAPSSLK","GLVSWGNIPC(UniMod_4)GSK","HVEPGEPLAPSPQEPQAVGR","VHKEDDGVPVIC(UniMod_4)QVEHPAVTGNLQTQR","NPDSSTTGPWC(UniMod_4)YTTDPTVR","QALPQVR"]

uniprot_list = list(df_protein["UniProt"].unique())
peptide_list = list(df_pebtide["Peptide"].unique())
patient_id_list = list(df_protein["patient_id"].unique())

#dropping skewed columns
peptide_list = [x for x in peptide_list if x not in ['ANRPFLVFIR', 'LQPLDFKENAEQSR', 'HQPQEFPTYVEPTNDEIC(UniMod_4)EAFRK']]
#the all and all pep may have error

selected_dic = {"selected":main_info_y_onehot+selected_pro+selected_pep}


model_params = {
    'layers': [[100, 50], [80, 50], [50,30] ,[30, 30], [100, 80, 80], [80, 80, 50], [80, 50, 50]],  # Number of neurons in each hidden layer
    'learning_rate': [1e-3],  # Learning rate for optimizer
    'dropout': [0.5],   # Dropout rate for regularization
}


updrs_list = ['updrs_1','updrs_2','updrs_3','updrs_4']

file_names = os.listdir("../dataset_cleaned_mk2/")

for output_prefix in selected_dic:
    selected_list = selected_dic[output_prefix]

    f_out = open("./20230331_"+output_prefix+"_training_result.csv","a")
    f_out.write("file_name,updrs,model_name,SMAPE,hyperparameter,run_time\n")
        
    for file_name in file_names:

        if "csv" not in file_name :
            continue
        df = pd.read_csv("../dataset_cleaned_mk2/"+file_name).astype('float32')
        df = df[selected_list]

        global custom_cv_dic 
        custom_cv_dic = get_cv_dictionary(df)

        for updrs in updrs_list:
            idx = list(df.columns).index("updrs_4")+1
            input_cols, output_cols = df.columns[idx:],  [updrs]


            st_time = time.time()
            best_params_,best_loss = train_model(df, input_cols, output_cols, model_params, list(custom_cv_dic.values()), max_epochs=200, batch_size=1200)
            took_time = str(time.time()-st_time)

            str_out = file_name+","+updrs+",DNN,"+str(float(best_loss))+ "," + str(best_params_).replace(",","|")+","+str(took_time)
            f_out.write(str_out+"\n")
            print(str_out)
```
