## Supplementary material (6) Additional Pipeline Codes for "Discovering Biomarker Proteins and Peptides for Parkinson’s Disease Prognosis Prediction with Machine Learning and Interpretability Methods": model_mk2_multivar.py

```
import warnings
warnings.filterwarnings('ignore')
import pandas as pd
import seaborn as sns
import os, code 
import numpy as np
import pickle
import matplotlib.pyplot as plt
import scipy.stats as stats
import time, random

from xgboost import XGBRegressor
from sklearn.ensemble import GradientBoostingRegressor
from sklearn.linear_model import BayesianRidge,LinearRegression,ElasticNet, SGDRegressor
from sklearn.tree import DecisionTreeRegressor

from sklearn.metrics import make_scorer
from sklearn.model_selection import GridSearchCV, ParameterGrid

from sklearn.model_selection import KFold
from sklearn.multioutput import MultiOutputRegressor

# Set random seed for reproducibility
seed = 42
random.seed(seed)
np.random.seed(seed)


def get_cv_dictionary(df):
	patient_ids = df['patient_id'].unique()
	kf = KFold(n_splits=5, shuffle= True, random_state=42)
	dic = {}
	for idx, (train_index, test_index) in enumerate(kf.split(patient_ids)):
		patient_train = patient_ids[train_index]
		patient_test = patient_ids[test_index]

		df_train_idx = np.where(df['patient_id'].isin(patient_train))[0]
		df_test_idx = np.where(df['patient_id'].isin(patient_test))[0]
		dic[idx] = (df_train_idx, df_test_idx)
	return dic


def weighted_mse(alpha = 1):
	def weighted_mse_fixed(label, pred):
		residual = (label - pred).astype("float")
		grad = np.where(residual>0, -2*alpha*residual, -2*residual)
		hess = np.where(residual>0, 2*alpha, 2.0)
		return grad, hess
	return weighted_mse_fixed


def run_training(f_out, df_y, df_X, updrs, file_name):
	models = [LinearRegression(),DecisionTreeRegressor(random_state=42),ElasticNet(random_state=42),BayesianRidge(),GradientBoostingRegressor(random_state=42)]
	#models = [models[-1]]
	def smape(y_true, y_pred):
		return np.mean((np.abs(y_true - y_pred) / (np.abs(y_true) + np.abs(y_pred)))) * 200

	# define the evaluation function to use SMAPE as scoring
	smape_scorer = make_scorer(smape, greater_is_better=False)

	param_grid = {
	 	"ElasticNet": {'alpha': [0.01, 0.1, 1.0],'l1_ratio': [0.1, 0.5, 0.9],'max_iter': [10000, 15000]},
		"DecisionTreeRegressor":{'max_depth': [2, 5, 10, 20, None],'min_samples_split': [2, 5, 10],'min_samples_leaf': [1, 2, 4],'max_features': [None, 'sqrt', 'log2']},
	 	"LinearRegression":{}}

	for model in models:
		st_time = time.time()
		modal_name = str(model).split("(")[0]

		grid_search = GridSearchCV(estimator=model, param_grid=param_grid[modal_name], cv=list(custom_cv_dic.values()), scoring=smape_scorer, n_jobs=16)

		# fit the grid search to the training data
		param_iterator = ParameterGrid(param_grid[modal_name])
		results = []
		for params in param_iterator:
			try:
				for train_idx, test_idx in list(custom_cv_dic.values()):
					# split data into train and test sets for cross-validation
					df_X = np.asarray(df_X)
					df_y = np.asarray(df_y)
					X_train, X_test = df_X[train_idx], df_X[test_idx]
					y_train, y_test = df_y[train_idx], df_y[test_idx]
					model.__init__(**params)
					# initialize gradient boosting regressor with current hyperparameters
					# fit model to training data
					model.fit(X_train, y_train)

					# make predictions on test data
					y_pred = model.predict(X_test) +0.00000001

					# calculate mean squared error on test data

					smape_tot = smape(y_test, y_pred)

					smapes = []
					for i in range(4):
						smapes.append(smape(y_test[:,i], y_pred[:,i]))
					smapes = [smape_tot] + smapes

					# store results as a tuple of hyperparameters and mean squared error
					results.append((str(params), "|".join([str(i) for i in smapes])))
			except:
				results.append(("error", "error"))
		# sort results by mean squared error
		results.sort(key=lambda x: x[1])

		# print best hyperparameters and mean squared error

		best_params_ =  results[0][0]
		best_score_ =  results[0][1]


		# The score may have a negative value due to the GridSearch pipeline problems, but can be replaced with positive numbers.
		took_time = time.time()- st_time
		str_out = file_name+","+updrs+","+modal_name+","+str(best_score_)+ "," + str(best_params_).replace(",","|")+","+str(took_time)
		f_out.write(str_out+"\n")
		print(str_out)


df_clinical = pd.read_csv("train_clinical_data.csv")
df_protein = pd.read_csv("train_proteins.csv")
df_pebtide = pd.read_csv("train_peptides.csv")


df_total = pd.read_csv("./dataset_cleaned_mk2/t_plus_0.csv")
df_total = df_total.drop(df_total.filter(regex='Unnamed').columns, axis=1)

main_info_y_onehot = list(['visit_id', 'patient_id', 'visit_month', 'updrs_1', 'updrs_2', 'updrs_3', 'updrs_4', 'upd23b_No', 'upd23b_Off', 'upd23b_On'])
selected_pro = ["O00533","O94919","P01009","P01019","P01857","P01860","P02787","P04156","P04433","P05060","P05452","P05546","P06454","P06727","P08603","P13591","P13987","P14618","P16035","P16152","P39060","P51884","P61916","P80748","Q06481","Q08380","Q12907","Q13332","Q92520","Q92823","Q96KN2","Q96PD5","Q99674","Q9BY67","Q9NQ79","Q99829"]
selected_pep = ["GNPEPTFSWTK","IEIPSSVQQVPTIIK","SMEQNGPGLEYR","VIAVNEVGR","VNGSPVDNHPFAGDVVFPR","AYQGVAAPFPK","ILEVVNQIQDEER","MGNFPWQVFTNIHGR","HSIFTPETNPR","NWGLGGHAFC(UniMod_4)R","DIPMNPMC(UniMod_4)IYR","LVDKFLEDVKK","ISLPESLK","TVAAPSVFIFPPSDEQLK","TPLGDTTHTC(UniMod_4)PR","DASGVTFTWTPSSGK","NFPPSQDASGDLYTTSSQLTLPATQC(UniMod_4)LAGK","VEPLRAELQEGAR","AKLEEQAQQIR","LEEQAQQIR","LGADMEDVC(UniMod_4)GR","LQAEAFQAR","VQAAVGTSAAPVPSDNH","WELALGR","GGSTSYGTGSETESPR","C(UniMod_4)SYTEDAQC(UniMod_4)IDGTIEVPK","VC(UniMod_4)PFAGILENGAVR","ITYGETGGNSPVQEFTVPGSK","NLQPASEYTVSLVAIKGNQESPK","RPGGEPSPEGTTGQSYNQYSQR","SYTITGLQPGTDYK","FSGTWYAMAK","SDVVYTDWKK","GSPAINVAVHVFRK","C(UniMod_4)C(UniMod_4)TESLVNR","KQTALVELVK","VHTEC(UniMod_4)C(UniMod_4)HGDLLEC(UniMod_4)ADDRADLAK","LC(UniMod_4)MAALK","DLLFRDDTVC(UniMod_4)LAK","IMNGEADAMSLDGGFVYIAGK","NFPSPVDAAFR","IYISGMAPR","LQSIGTENTEENRR","VVEQMC(UniMod_4)ITQYER","LEPGQQEEYYR","SSGLVSNAPGVQIR","GYPGVQAPEDLEWER","SSQGGSLPSEEK","ISYGNDALMPSLTETK","STNLHDYGMLLPC(UniMod_4)GIDK","WYFDVTEGK","C(UniMod_4)PNPPVQENFDVNK","LLDSLPSDTR","NSVIKVPMMNSK","C(UniMod_4)RDQLPYIC(UniMod_4)QFGIV","AGALNSNDAFVLK","AGKEPGLQIWR","EVVEEAENGR","VSTLPAITLK","C(UniMod_4)KPVNTFVHEPLVDVQNVC(UniMod_4)FQEK","VTGVVLFR","NANTFISPQQR","EDC(UniMod_4)NELPPRR","RLEAGDHPVELLAR","C(UniMod_4)FSGQC(UniMod_4)ISK","GLSAEPGWQAK","RLEGQEEEEDNRDSSMK","YPGPQAEGDSEGLSQGLVDREK","KTLLSNLEEAK","TLLSNLEEAK","TLLSNLEEAKK","LRENELTYYC(UniMod_4)C(UniMod_4)KK","AVSEKEVDSGNDIYGNPIKR","SC(UniMod_4)SPELQQK","SNAQGIDLNR","SSALDMENFR","MVQEQC(UniMod_4)C(UniMod_4)HSQLEELHC(UniMod_4)ATGISLANEQDR","RGYQLSDVDGVTC(UniMod_4)EDIDEC(UniMod_4)ALPTGGHIC(UniMod_4)SYR","QKWEAEPVYVQR","LQDLYSIVR","FSGSNSGNTATLTISR","AAFGQGSGPIMLDEVQC(UniMod_4)TGTEASLADC(UniMod_4)K","NLHGDGIALWYTR","SSNTYTLTDVRR","SRYPVC(UniMod_4)GSDGTTYPSGC(UniMod_4)QLR","LVWEEAMSR","LIADLGSTSITNLGFR","VFNTPEGVPSAPSSLK","GLVSWGNIPC(UniMod_4)GSK","HVEPGEPLAPSPQEPQAVGR","VHKEDDGVPVIC(UniMod_4)QVEHPAVTGNLQTQR","NPDSSTTGPWC(UniMod_4)YTTDPTVR","QALPQVR"]
uniprot_list = list(df_protein["UniProt"].unique())
peptide_list = list(df_pebtide["Peptide"].unique())
patient_id_list = list(df_protein["patient_id"].unique())

#dropping skewed columns
peptide_list = [x for x in peptide_list if x not in ['ANRPFLVFIR', 'LQPLDFKENAEQSR', 'HQPQEFPTYVEPTNDEIC(UniMod_4)EAFRK']]


#re run the all peps
selected_dic = {
"selected":main_info_y_onehot+selected_pro+selected_pep}

for output_prefix in selected_dic:
	selected_list = selected_dic[output_prefix]
	updrs_list = ['updrs_1','updrs_2','updrs_3','updrs_4']
	f_out = open("./results_mk2/multivar/20230402_"+output_prefix+"_training_result_dummy.csv","a")
	f_out.write("file_name,updrs,model_name,SMAPE,hyperparameter,run_time\n")
	for dataset_path in os.listdir("dataset_cleaned_mk2"):
		print(dataset_path)
		if ".csv" not in dataset_path:
			continue
		df = pd.read_csv("./dataset_cleaned_mk2/"+dataset_path)[selected_list]
		global custom_cv_dic 
		custom_cv_dic = get_cv_dictionary(df)

		for updrs in updrs_list:
			idx = list(df.columns).index("updrs_4")+1
			df_y = df[updrs_list]
			df_X = df[df.columns[idx:]]
			run_training(f_out,df_y, df_X,updrs, dataset_path)
		
```
